# A mathematical model of photoinhibition: exploring the impact of quenching processes

**DOI:** 10.1101/2023.09.12.557336

**Authors:** Tim Nies, Shizue Matsubara, Oliver Ebenhöh

## Abstract

Plants are constantly exposed to changing environments, sometimes leading to extreme conditions and stress. For example, sudden exposure to high light leads to excess absorbed light energy, causing reactive oxygen species (ROS) formation. ROS damage the photosynthetic machinery, particularly the D1 protein in photosystem II (PSII), which therefore needs to be continuously repaired and replaced. The effect of the damage inflicted by high light is a prolonged decrease in photosynthetic efficiency. Hence, it is not surprising that photoinhibition has been subject to numerous experimental studies investigating its effects in the context of crop productivity. However, it has become apparent that classical measures of photoinhibition, i.e., changes in the chlorophyll fluorescence parameter F_v_/F_m_, are not only determined by the loss of PSII core function but also by processes such as energy transfer and quenching. Mathematical models can help dissect the influences on such fluorescence signals and quantify the contributions of various interacting mechanisms. We present a mathematical model with a dynamic description of the photosynthetic electron transport chain (PETC), non-photochemical quenching, and photoinhibition. With our model, we investigate the interconnection between quenching, photoprotection, and fluorescence using simulations and experimental data. We found that different energy-dissipating properties of intact and damaged PSIIs, as well as energy transfer between PSIIs, are critical components that need to be included in the model to ensure a satisfactory fit to the experimental data. We envisage that our model provides a framework for future investigations of photoinhibition dynamics and its importance for plant growth and yield.

## Introduction

Photosynthesis is one of the main processes that make energy available to the biosphere [16]. By capturing light, photosynthetic organisms convert solar energy into usable chemical energy, which is then used to drive metabolic processes, including the formation of biomass. Plants, algae, and other photosynthetic organisms exist in a wide range of environments, ranging from deserts to tropical forests. These environments can exhibit drastically and rapidly changing external conditions, considering, e.g. light intensity, temperature, and humidity. Plants, as sessile organisms, must adapt to the conditions they are exposed to [11]. However, such fluctuating conditions make the coordination of the photosynthetic electron transport chain (PETC), supplying light energy in the form of ATP and NADPH, and the Calvin Benson Bassham cycle (CBB cycle), which uses ATP and NADPH to sequester CO_2_, a challenging task [28]. Antenna complexes in chloroplast thylakoid membranes collect light energy and channel it to the reaction centers of the PETC. This captured energy is used to drive photochemistry, but the excited states can also dissipate energy as heat or re-emit it as fluorescence [26]. Due to variations in external conditions, the light energy supply can frequently exceed the demand, which leads to the formation of reactive oxygen species (ROS) at multiple sites of the PETC. ROS are highly reactive compounds that can damage the molecular machinery of the PETC [12].

The photodamage induced by ROS affects various proteins, with the D1 subunit of photosystem II (PSII) being the most susceptible. In fact, with a turnover rate of *>* 0.5 d^*−*1^, the D1 subunit exhibits one of the shortest protein lifetimes in the PETC [18]. For functional photosynthesis, it is therefore essential that this protein is constantly resynthesized and replaced. This is realized by the so-called D1 protein repair cycle, which involves the degradation and synthesis of damaged D1 protein. This cycle has a very high energy demand, with an estimated 1304 ATP per subunit repaired [27]. Despite considerable advances in our understanding of photoinhibition, the exact mechanism of how high-light stress inflicts damage on the photosynthetic machinery is still under debate, and various hypotheses have been proposed [45].

Classically, photoinhibition is quantified by measuring F_v_/F_m_ after prolonged exposure to strong irradiance. This was justified because of the almost linear relationship between F_v_/F_m_ and the loss of photosynthetic O_2_ evolution (see, e.g. [31]). It has recently become increasingly apparent that the F_v_/F_m_, derived from the fluorescence signal, might not be ideal for assessing photoinhibition. The fluorescence signal that a photosynthetic tissue, such as a leaf, emits is influenced by multiple factors, such as nonphotochemical quenching, the efficiency of photochemistry, and the three-dimensional structure of the leaf. Hence, F_v_/F_m_ might be determined not only by the loss of the PSII core function but also by other dissipating processes [20]. Moreover, also theoretical studies have suggested an inherently nonlinear relationship between inactive PSII and the fluorescence signal [7].

Over the last decades, various mathematical models of photosynthesis were developed [39]. Some of them focus on the PETC [6, 22, 44] or the CBB cycle [32, 33], and others try to integrate both into one mathematical description [25, 23, 36]. Other models focused on detailed processes in PSII [2]. Many of these models calculate how the fluorescence signal derives from the molecular processes of the PETC. Most of the calculations depend on equations that describe the fluorescence yield associated with closed and open reaction centers of PSII. The difference in how these models determine fluorescence yield primarily arises from different simplified or extended versions of these equations. These equations are based on the current understanding regarding the source of the fluorescence signal, derived from the work conducted during the last sixty years [10, 5, 14, 7, 3]. However, despite much effort, it still needs to be clarified which of the classical equations and which model representation of the thylakoid membrane (e.g., lake, single unit, domain model, see [3]) is most realistic.

Here we expanded a published model of the PETC and non-photochemical quenching (NPQ) [6, 22, 36] by integrating a mechanistic description of photoinhibition and the D1 repair cycle. For this, we build upon previous models of the D1 damage-repair cycle and an expansion of the energy transfer theory [7, 40, 31]. The goal of our model is to quantitatively reproduce experimental data measuring photodamage as changes in F_v_/F_m_, F_m_, and F_o_ in wildtype *Arabidopsis thaliana* and the *npq1* mutant. The *npq1* mutant lacks violaxanthin de-epoxidase and, thus, zeaxanthin. Zeaxanthin has been shown to play a critical role in the induction of short(qE) and longterm (qZ) quenching processes, potentially protecting against high light-induced damage [9, 29]. Our model provides a theoretical framework in which we discuss different formulations for the fluorescence yield based on previous work and assess how these agree with experimental data. In particular, we focus on the effects of different heat dissipation capabilities and quenching activities on the fluorescence signal under photoinhibtory conditions. This work helps to clarify which processes contribute to the dynamic changes of photosynthesis under high-light stress. Moreover, we provide a quantitative and mechanistic explanation of the observed changes in F_v_/F_m_, F_o_, and F_m_ during high light-induced photoinhibtion.

## Results

For our analysis, we constructed a mathematical model that combines the description of the PETC as in [6, 22] and the D1 damage-repair cycle from [40] (for details, see Methods and Supplement). In the following, we describe the development of hypotheses about mechanistic aspects of the fluorescence signal during photoinhibition and compare model predictions with experimental observations. Guided by discrepancies between experiment and simulations, we iteratively refine our hypotheses to arrive at a realistic description of the fluorescence signal.

### Experimental dynamics of fluorescence signals

The data (see Fig. S1) comprises F_v_/F_m_, F_m_ and F_o_ measurements for *Arabidopsis thaliana* wildtype and *npq1* mutant plants for different exposure times to high light and with or without treatment with lincomycin, which inhibits chloroplast protein synthesis and thus the D1 repair (see Methods). The experimental data suggest that the *npq1* mutant, which lacks violaxanthin de-epoxidase enzyme and thus cannot form zeaxanthin in the so-called xanthophyll cycle, reacts more sensitively to high-light stress in water (control) and lincomycin treatment. Fig. S1 shows that the relative reduction of F_m_ is generally more pronounced than the increase of F_o_, indicating F_m_ to be the main factor determining the changes in F_v_/F_m_ in this experiment. While the differences between the water and lincomycin treatment are clearly discernible for the wildtype and *npq1* mutant in the F_m_ and F_v_/F_m_ signal, this is not the case for F_o_.

### Changes in the F_v_/F_m_ signal

We started our computational analysis with the most simple assumptions for the model extended with photoinhibition: We assume that 1) the duration and intensity of the high-light treatment determine the amount of inactive PSII; 2) inactive PSII contributes to fluorescence and has the same quenching properties as active PSII and; 3) there is no energy transfer between active and inactive reaction centers. With these assumptions, our model of photoinhibtion cannot reproduce the experimentally observed data (see Fig. S1). The increase of F_o_ with prolonged highlight treatment is much higher than in the experiment, while there is only little or no effect for simulated F_m_. Interestingly, the F_v_/F_m_ signal can be described by the model, indicating that the F_v_/F_m_ signal alone does not provide sufficient information to understand the underlying mechanisms.

### Fluorescence signal in photoinhibtion

Motivated by this observation, we modified our model similar to [7] by assuming that the fluorescence signal and heat dissipation properties of active and inactive PSII can differ. This means we relax assumption 2 stated above. To quantify the different behaviour, we introduce the parameter *ρ* as the ratio of heat dissipation rate constants between inactive and active states of PSII – see Eq.(6). This means that *ρ* = 1 corresponds to the previous model, *ρ <* 1 denotes a model in which inactive PSII dissipate heat less effectively and thus yield more fluorescence than active PSII, and *ρ >* 1 describes the opposite scenario.

Using Eqs. (7) and (9), we can predict the qualitative changes of *F*_*m*_ and *F*_*o*_ as a response to photodamage:

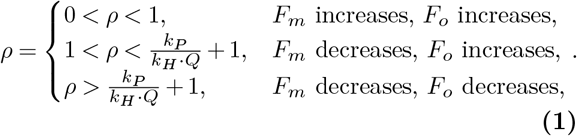

An increase or decrease of F_m_ depends only on whether *ρ* is larger or smaller than 1. In contrast, the *F*_*o*_ behavior (increase or decrease) depends not only on the value of *ρ* but also on the quenching activity *Q*.

Fig. 2 shows that for the case in which the heat dissipation of active and inactive photosystems is identical (*ρ* =1, continuous lines), F_v_/F_m_ follows a linear relationship with the fraction of active PSII, both in a low and high quenching scenario (black, and red lines). However, the relationship becomes nonlinear when the active and inactive PSII differ in their heat dissipation capabilities. We further observe that *ρ* determines the curvature of the relationship be224 tween F_v_/F_m_ and active PSII fraction, while an active quencher makes the non-linearity more pronounced. The dependence of *F*_*m*_ and *F*_*o*_ on active PSII is linear in all cases. However, the slope is affected both by *ρ* and *Q*. Note that *F*_*m*_ is not affected by photoinhibition for *ρ* = 1 (original model, Fig. S1, see also [7]).

**Figure 1.**
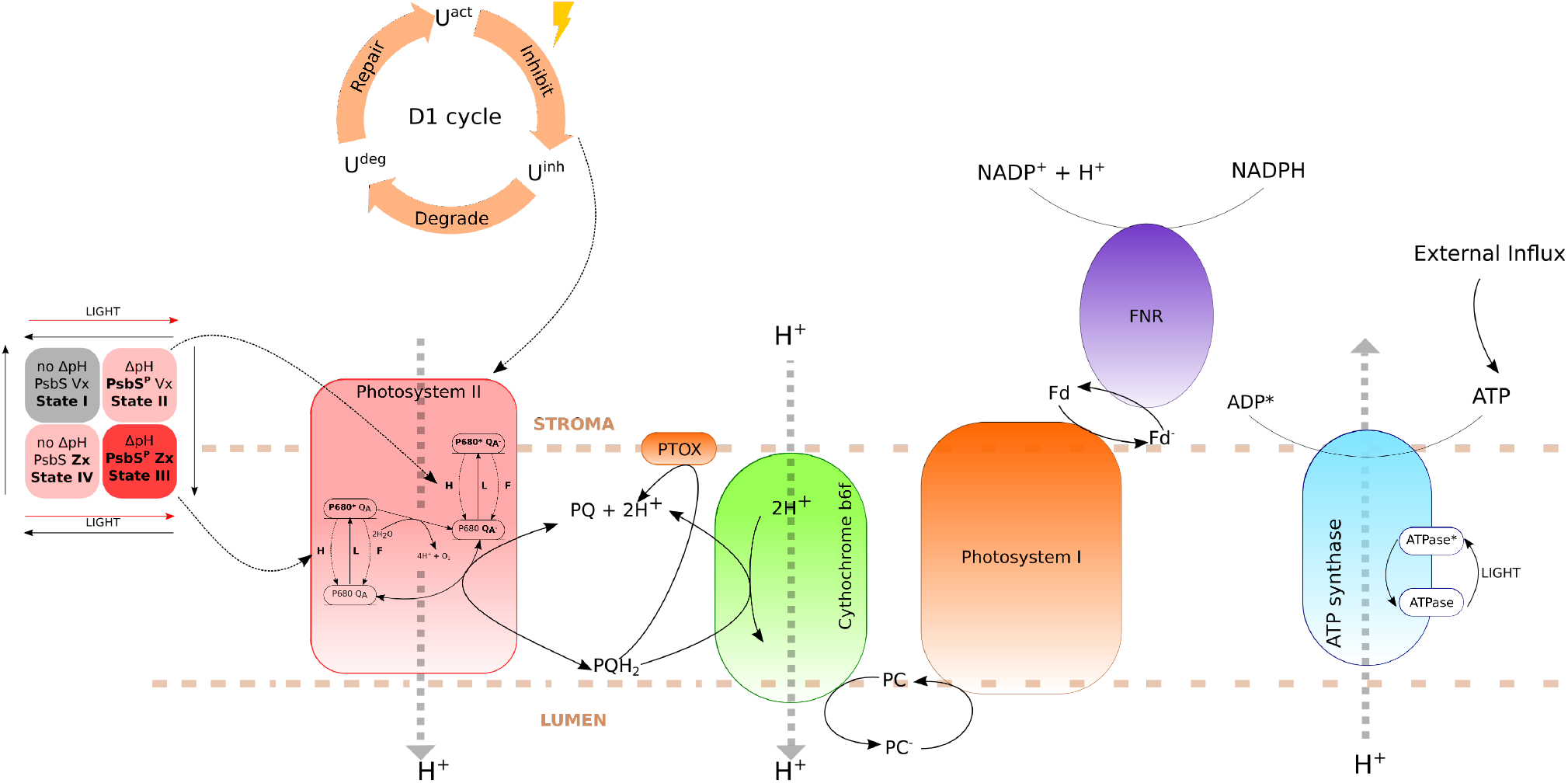
Schematic depiction of the model of photoinhibition (compare also [6, 22]). Not shown for clarity but included are the cyclic electron flows around photosystem I.

**Figure 2.**
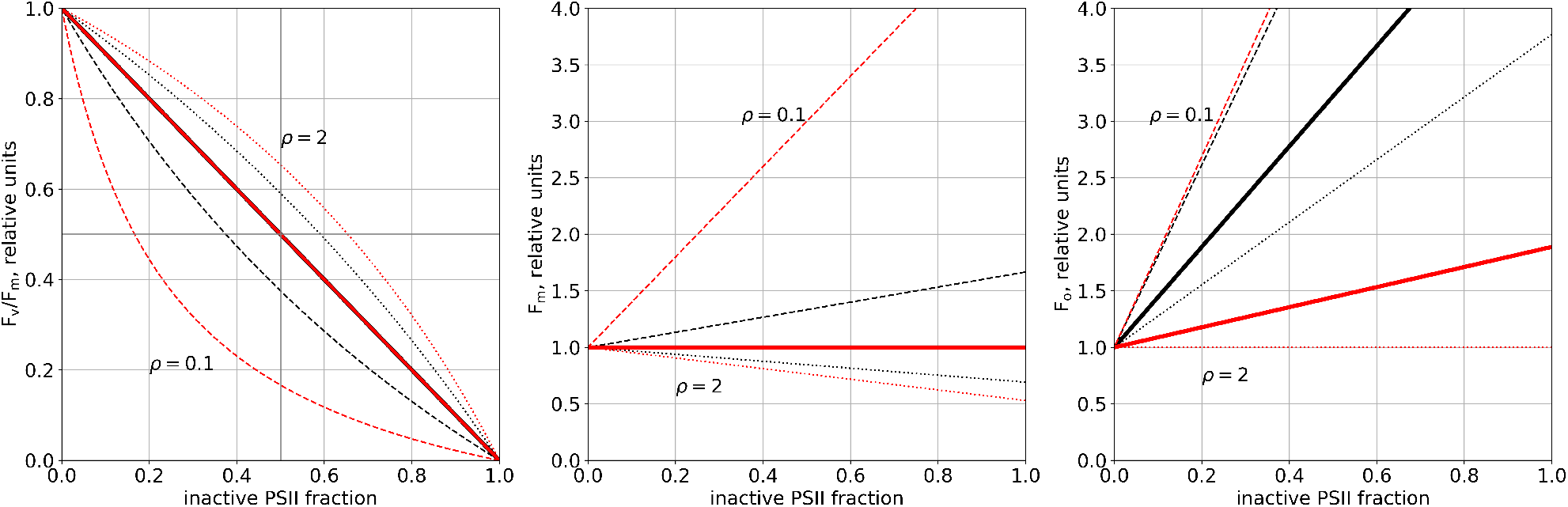
Relationship between F_v_/F_m_, F_m_, and F_o_ and fraction of inactive photosystem II based on Eq. 6. The dashed, continuous, and dotted lines indicate scenarios in which the ratio of heat dissipation between inactive and active PSII is 0.1, 1, and 2, respectively. Black lines signify a low quenching, while red lines denote high quenching activity (Q = 0.1 and 1). Parameter values used for the calculations can be found in the supplement table S1

For a fluorescence yield model without energy transfer, the ratio of the slopes of relative values of F_o_ and F_m_ as functions of active PSII is given by Eq. (13). The slope ratio has a singularity at *ρ* = 1 where the slope of F_m_ becomes zero. The slope ratio is zero at *ρ* = *k*_*P*_ */*(*k*_*H*_ *··Q*) + 1, when the slope of F_o_ is zero. In our fluorescence measurements for *A. thaliana* during high-light treatment, we observed that the realtive increase of F_o_ is smaller than the relative decrease of F_m_. To reproduce this behavior, the slope ratio must be negative, in the range between -1 and 0. For this, *ρ* must be constrained to the interval

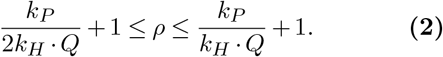

Fig. 3 depicts the slope ratio for the parameter values in the model for two different quenching activities. In a low quenching scenario (*Q* = 0.1, solid line), the parameter *ρ* is predicted to lie in the range between 6 and 11. This means that, in order to reproduce the experimentally observed slope ratio, damaged PSII needs to dissipate heat with a rate at least six times larger than that at which intact PSII does. Similarly, in a high quenching scenario (*Q* = 1, dotted line) we find 1.5 *≤ ρ≤* 2, which means a oneto twofold faster heat dissipation for damaged vs. active PSII.

**Figure 3.**
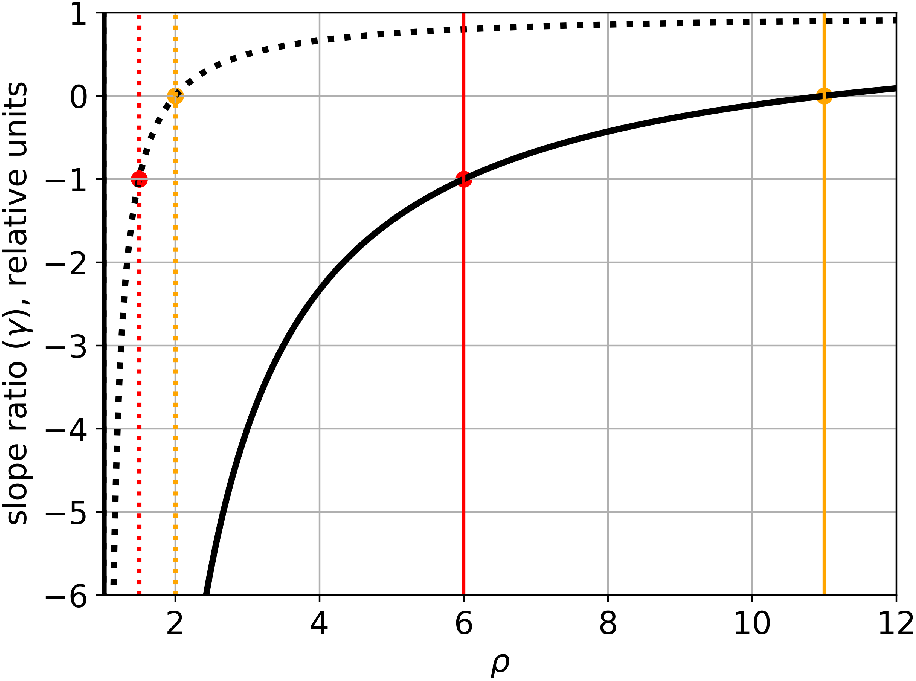
The slope ratio *γ* for model without energy transfer in a high (dotted line) and low quenching scenario (continuous line). Vertical lines indicate the points at which the slope ratio is -1 or 0. Parameters are the same as for Fig. 2.

We used these constraints to fit our model to the experimental data. We find that the data could be considerably better explained than in the model with *ρ* = 1 (see Suppplementary Figs. S2 and S3). With the parameter *ρ* in the range determined above, all qualitative features of the fluorescence traces could be reproduced. However, there are still quantitative discrepancies, which could not be resolved using this model.

We therefore expanded the model to include excitation energy transfer between closed active and damaged PSII, following the example in [7]. This leads to a modified formula to describe F_m_, whereas the description for F_o_ remains the same as in the case without energy transfer (see Eqs. 17 and 18). Consequently, the relation between F_m_ and active PSII becomes nonlinear (see Fig. 4). The effect of an excitation energy transfer between active and inactive PSII leads to a faster decrease of the F_v_/F_m_ value in response to lowering the active PSII fraction. Moreover, the effect of the energy transfer seems to be larger in a low quenching than a high quenching state (compare Figs. 2 and 4). Because the description of F_o_ does not change compared to the isolated case, *ρ* and the quencher activity are still the determining factor for the behavior of F_o_. However, the behavior of F_m_ is a nonlinear function of the active PSII fraction, and therefore a slope ratio can no longer be uniquely defined.

**Figure 4.**
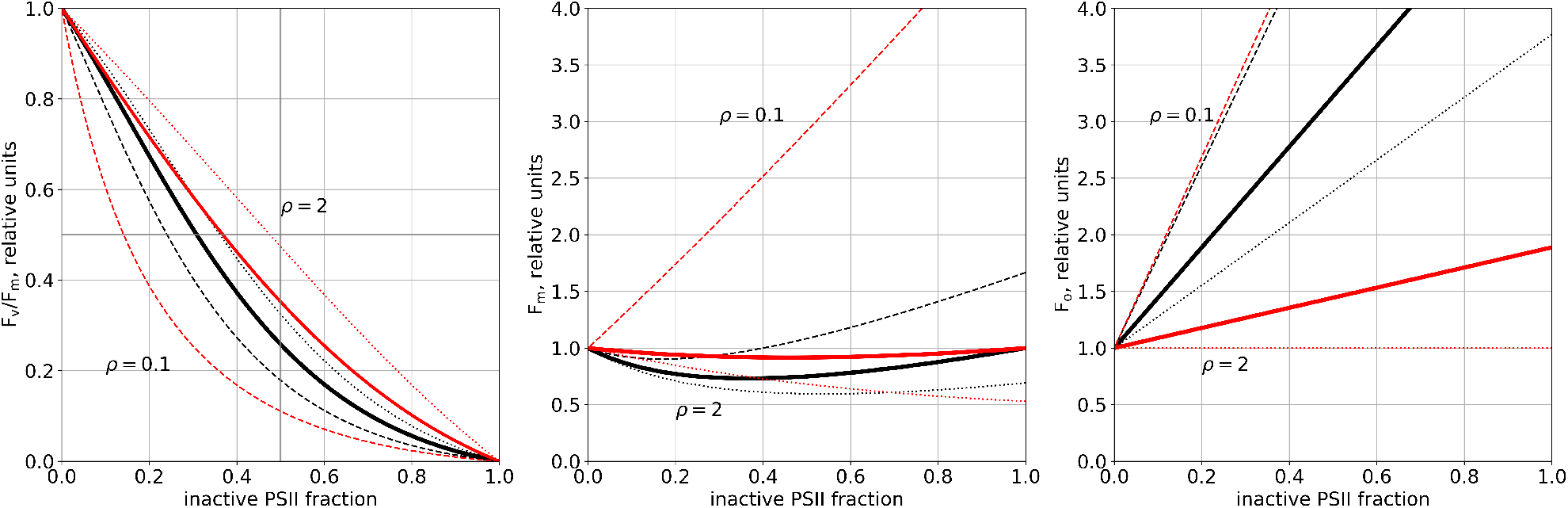
Relationship between F_v_/F_m_, F_m_, and F_o_ and fraction of inactive photosystem II based on Eq. 16. The dashed, continuous, and dotted lines indicate scenarios in which the ratio of heat dissipation between inactive and active PSII is 0.1, 1, and 2, respectively. Black lines signify a low quenching, while red lines denote high quenching activity (Q = 0.1 and 1). Parameter values used for the calculations can be found in the supplement table S1. Energy transfer was set to 8 10^8^ mmol^*−*1^ (mol Chl) s^*−*1^.

### Model predictions

Guided by comparison of model predictions and experimental data, we have iteratively refined a model of the photosynthetic electron transport chain. The resulting model includes the assumption that energy quenching differs between active and damaged photosystems. Morever, energy can be transferred from active to damaged photosystems. This model version can satisfactorily reproduce our experimental data for *A. thaliana* (see Fig. 5). In the following, we employ our model to make novel predictions how photoinhibition affects key photosynthetic parameters.

**Figure 5.**
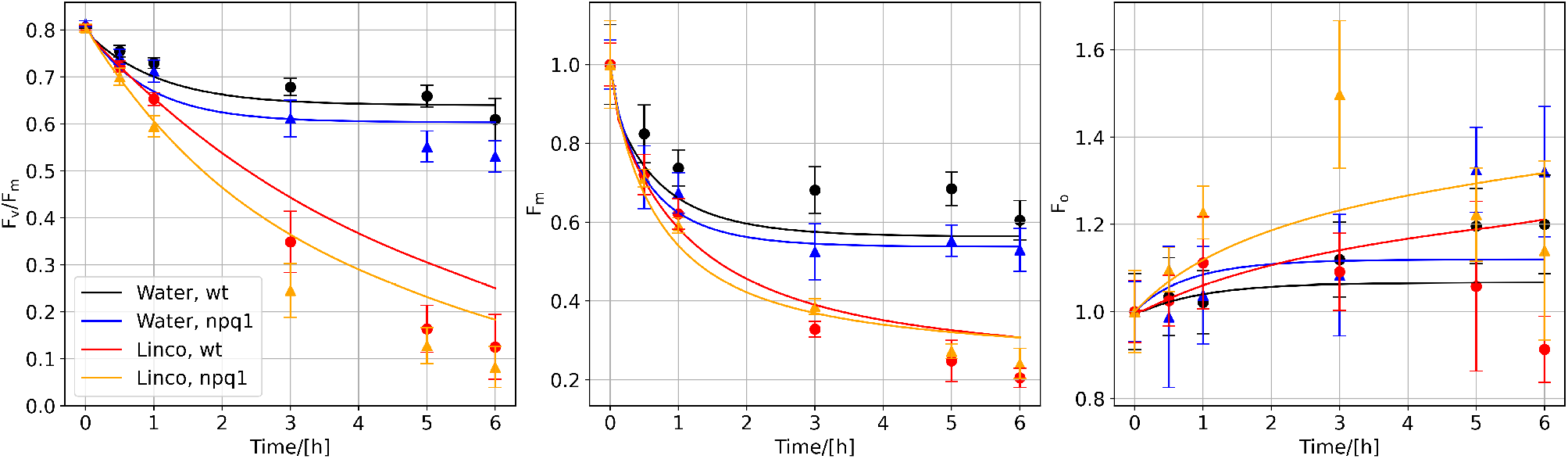
Experimental measurement and simulated changes in F_v_/F_m_, F_m_, and F_o_ in high-light treatment of *A*.*thaliana* plants for 6 hours. The plants were either treated with water (black and blue lines) or lincomycin (red and orange line) inhibiting protein synthesis. Light intensity was 800 µmol m^−2^ s^−1^.

#### Quenching shifts the fraction of closed and open PSII during photoinhibition

To describe internal processes of photosystem II, we used a simplified mathematical representation that has been applied successfully for modeling fluorescence signal changes in connection to state transition and non-photochemical quenching [6, 22, 23]. This representation of PSII can be approximated by a two-state system consisting of the open and closed active PSII states.

Fig. 6 shows the changes of closed and open active PSII states during exposure to various light intensities for four hours as phase-space trajectories. We investigate four model versions with (right column) and without (left column) dynamic quencher activity as well as with non-constantly (top row) and constantly active (bottom row) ATP synthase. The version with non-constantly active ATP synthase and dynamic quencher is our original model (top left). For all four versions, the phase-space provides information about the different stages we observe during the onset of photoinhibition. These stages are characterised by the different time-scales on which they operate. The simulation starts with a dark-adapted state and, hence, with no closed PSII. When the light is switched on, the system almost instantaneously changes to a state where both closed, and open PSII are present. The ratio of open to closed PSII depends on the light intensity. A light intensity of around 1000 µmol m^−2^ s^−1^results in approximately 85% of PSII in the closed state. This initial stage is driven by the rapid processes in photosystem II.

**Figure 6.**
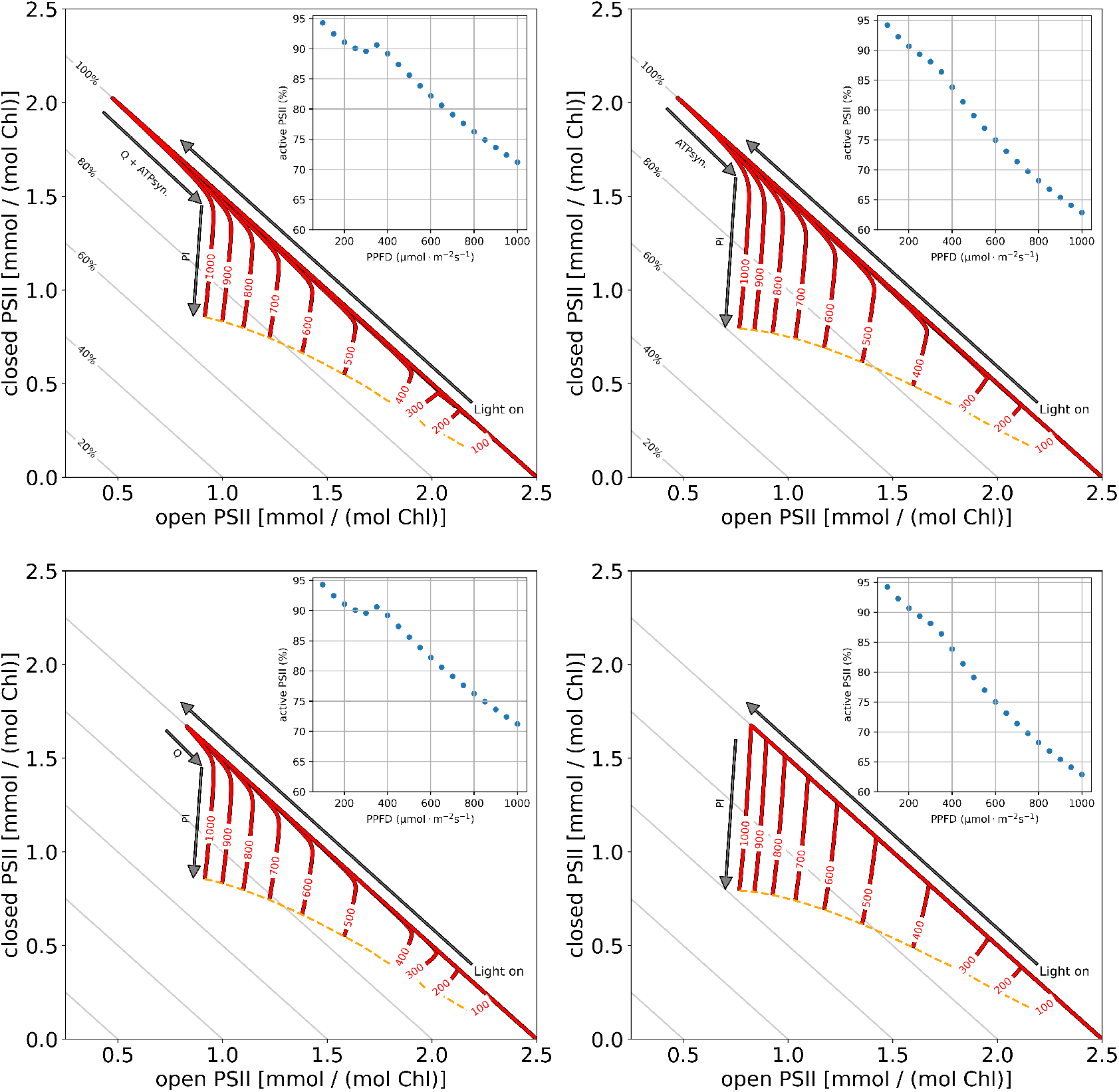
phase-space of open (B0) and closed (B2) active PSII states during photoinhibtory treatment in various light intensities (100 — 1000 µmol m^−2^ s^−1^). Red lines indicate changes in open and closed PSII. The orange dashed line connects all points in the phase-space reached after 4 hours of light treatment. Grey lines indicate the fraction of total active PSII. Inset shows the fraction of active PSII as a function of applied light intensity at the end of the simulation. The top left and top right panel show the phase-space of a model version with and without a dynamic quencher. The bottom left and bottom right show the phase-space of a model version with and without a dynamic quencher and without ATP synthase activation.

The first stage is followed by the second stage, which operates on a time-scale of seconds to minutes. In this phase, two effects dominate. Firstly, ATP synthase is activated (arrows marked as “Q + ATPsyn.” and “ATPsyn”.). Secondly, the fast component of the quencher is rapidly activated, leading to a slower activation of PSII and thus a smaller fraction of closed states (compare top row with bottom row). Comparing the left (dyanmic quencher) and right (no quencher) columns as well as the top (non-constantly active ATP synthase) and bottom (constantly active ATP synthase) rows of Fig. 6 illustrates the effect of these two processes individually. In this stage, photoinhibition starts to become active but photodamage is still negligible.

This stage is followed by the slower stage of photoinhibition, which extends over several hours. Here, the active amount of PSII is gradually reduced due to photodamage. In the phase-space this is reflected by the downward pointing red lines. This phase continues until repair processes compensate for the extent of the light-induced damage, indicated by the dashed yellow lines. By comparing the four model versions with and without a dynamic quencher and with non-constantly and constantly active ATP synthase, it becomes apparent that quenching not only leads to more open PSII but also reduces the extent of photodamage, visible by the shorter downward trajectories for the model with active quencher. In our model simulation and with our chosen parameters, the quenching activity leads to almost 10% more active PSII after four hours of light treatment with an intensity of 1000 µmol m^−2^ s^−1^(see inset in Fig. 6).

#### Steady state photoinhibition analysis

We observed that dynamic quenching, associated with PsbS and the xanthophyll cycle (Fig. 5), is a key determinant for the extent of high-light stress-induced photodamage. We employed our model to systematically analyze the connection between quenching and the steady-state behavior for different light intensities. For this, rate constants associated with non-photochemical quenching were set to zero, and the quenching activity was fixed to be a constant value. Subsequently, the system was simulated until it reached a steady state. Fig. 7 displays the computed steady state photoinhibition rate.

**Figure 7.**
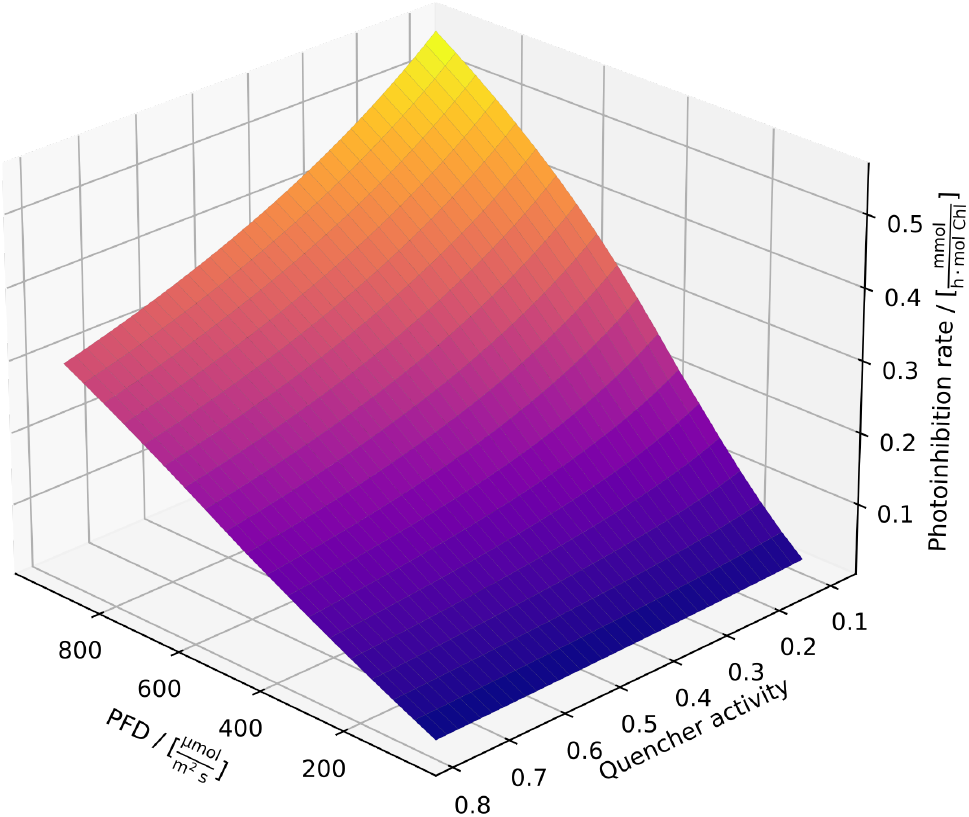
The predicted stationary flux of photoinhibition for different light intensities and different quencher activities. Quenching activities were modeled for these predictions by imposing fixed values (x-axis) between 0.1, representing almost no quencher activity, and 0.8., representing double the quencher activity typically observed in our model simulations. Light intensity (x-axis) was varied. The system was simulated for each combination of light intensity and quenching activity until a steady state was reached. On the z-axis, the stationary photoinhibition rate is plotted. For low quenching activities, a sigmoidal transition between high and low photoinhibition rates is observed with light intensities. This demonstrates that small light intensity changes can already have strong photoinhibitory effects in low light regimes. For higher quenching activities, the transition is smoother, almost linear, indicating higher flexibility, as expected, more flexibility against high-light stress.

In low quenching regimes, we observe a slightly sigmoidal transition between high and low photoinhibition rates with increasing light intensities. For very low quenching activities, the photoinhibition rate increases quickly, having a disproportionally high increase at around 400 µmol m^−2^ s^−1^. This demonstrates that small light intensity changes can already have strong photoinhibitory effects in low light regimes. By contrast, when quenching is active, we observe a smooth transition from low to high-light intensities, indicating greater tolerance against highlight stress.

## Discussion & Conclusions

We have presented a model of the PETC integrating non-photochemical quenching and photoinhibitory processes. The model aims to a) investigate how fluorescence signals (F_m_ and F_o_) in response to photoinhibition can be explained, b) explore which assumptions are sufficient to reproduce experimental data, c) study the effects of different modes of energy quenching, and d) quantify stationary photoinhibtory rates. To do so, we followed a reductionist approach. Our initial model version of photodamage in the PETC was built on the simple assumptions that 1) photoinhibition is proportional to intensity and duration of light treatment, 2) there is no difference between heat dissipation properties of active and damaged photosystems, and 3) there is no energy transfer between photosystems. However, this version could not reproduce the experimental data; see Fig. S1. Motivated by differences between simulations and experimental data, we systematically increased the complexity of the model representation by firstly introducing differences in heat dissipation properties of active and inactive photosystems (Fig. 2) and secondly an energy transfer between closed active and inactive photosystems in the description of the fluorescence signal (Fig. 4).

Both additions are realistic and have previously been used to study fluorescence changes after high-light treatment [7] with fluorescence yield models. This previous investigation did not include a dynamic lumenal pH-induced quenching component. Our model implements quenching based on the four state model introduced by [9], which is also included in [22]. In comparison to [22], in our model version the influence of the qZ component (zeaxanthin concentration high, no PsbS protonation) is reduced. This modification was necessary to realistically simulate differences between the wildtype and *npq1* mutant. After introducing all these changes to our initial model version, the fit agreed with experimental data (Fig. 5), supporting the assumptions that heat dissipation properties differ between intact and damaged photosystems and that energy transfer occurs.

Whereas the F_v_/F_m_ and F_m_ signals could be very well reproduced with deviations in the range of experimental errors, the experimental F_o_ signal slightly deviates from our simulated fluorescence traces. We hypothesized that a long-term quencher independent of the xanthophyll cycle and PsbS protonation, not yet implemented, could improve the model fit. To test this, we implemented an additional component in the quencher description of [22] proportional to inactive PSII to our first model version (without heat dissipation differences and energy transfer). This should mimic a quenching process proportional to high-light stress that is still strongly active after dark adaption. Although the changes in the F_m_, F_v_/F_m_ and F_o_ signal are now primarily products of the long-term quencher (compare Fig. S1 and Fig. S4), the agreement between simulated and experimental fluorescence traces improved, even reproducing the decrease of the F_o_ signal in lincomycin treatment (Fig. S4). However, the conditions under which we recorded the experimental data should not induce any additional strong longterm quenching component, motivating us to discard this long-term quencher hypothesis and instead to focus on the initial simple description of fluorescence yield based on [7].

Besides replicating experimental data, the value of a model lies in providing a way to investigate biological phenomena not easily accessible by experiments. Here, we specifically focused on the changes in excited and non-excited active PSII during photoinhibition. We used a phase-space visualization to observe the dynamic response of the system to different light conditions (Fig. 6). Our results show that one effect of the quencher is to actively push PSII to more open states, leading to a long-term reduction of high-light induced photodamage. The changes in active PSII shown in Fig. 6 is probably due the activation of the ATP synthase and the saturation of the quencher. Additionally, we investigated the effects of quenching for steady-state rates of photodamage and found a disproportionally strong effect of highlight stress in low-quenching scenarios (Fig. 7). In high-quenching scenarios, the response becomes linear, indicating that quenching might be essential for the flexible behavior of photosynthetic organisms under high-light stress.

Combining the previous observations, we might speculate that fluorescence changes induced by high-light stress are caused by a combination of various processes, including the reduction of PSII core functionality and multiple long- and short-term quenching mechanisms. Our simulations indicate that, to explain observed changes in the F_v_/F_m_, F_m_ and F_o_ signals, three components are essential: 1) the amount of active and inactive PSII, 2) the difference between their heat dissipation properties and 3) quenching phenomena. For the latter, it is essential to distinguish between shortand long-lived quencher components. While short-lived quenchers influence the decrease of the active PSII fraction but not the fluorescence signal measured after dark-adaption, long-lived quenchers influence both.

There is a continuous discussion about whether inactive PSII is photoprotective [21, 37, 15]. This hypothesis was based on the observation that an active PSII pool remained even after prolonged high-light treatment and repair inhibited by lincomycin [17]. However, later studies did not support these findings and it was speculated that the observed active pools resulted from the specific experimental setup [15]. Regarding the mechanism, it was hypothesized that photoprotection is caused by an energy transfer from active to inactive photosystems, which are more efficient energy quenchers [21]. It was argued that without energy transfer photoinhibition is a first-order process, and that the existence of an energy transfer and photoprotection should be detectable by a deviation from an exponential kinetics [21, 37].

With our model, we can test these hypotheses by simulating the respective scenarios. Fig. S5 shows the dynamics of PSII simulated with (red) and without (orange) assumed energy transfer. We observe that in both cases the dynamics of active PSII closely resemble a simple exponential, and thus may be interpreted as a first-order process. However, even in the case without energy transfer, small discrepancies from the exponential behavior are visible. Although such small differences are unlikely to be experimentally detectable, they can be theoretically explained. An exact exponential decay would entail that the fraction of excited PSII (relative to active PSII) remains constant. However, in our simulations this is not precisely the case (see Fig. S6). The cause for this is that the redox state of the plastoquinone pool and the state of the quencher depend on the rate of electrons provided by PSII, and thus on the amount of active PSII itself, leading to a non-trivial dynamics which is only approximately exponential. Interestingly, even the decay of PSII under the assumption of energy transfer closely resembles an exponential. We therefore conclude that observing discrepancies from an exponential behaviour might not be the best suited method to discriminate between the two hypotheses.

This is especially the case when using F_v_/F_m_ as a measure of photoinhibtion. Our calculations have shown that, in a scenario without energy transfer, changes in F_v_/F_m_ only follow the active PSII decay proportionally if the active and inactive PSII have identical heat dissipation properties (*ρ* = 1, see Fig. 2). However, because we used F_m_ and F_o_, besides F_v_/F_m_, to guide our simulations, we could show that the experimental observations can only be explained if *ρ >* 1, which means that inactive PSII quench energy more efficiently than active PSII. This in turn means that F_v_/F_m_ is a nonlinear function of inactive PSII, and as a consequence the F_v_/F_m_ signal displays a slightly different kinetic than the active PSII pool (see Figs. S5 and S7). Nonetheless, without energy transfer also a value of *ρ >* 1 results in simulated F_v_/F_m_ that is too large compared to the experiment (see Figs. S2 and S3). Assuming an energy transfer, leads to reduced simulated F_v_/F_m_ values and allows quantitative reproduction of the measured signal (Figs. 4 and 5). Interestingly, energy transfer leads to a more linear response of the F_v_/F_m_ signal to inactive/active PSII (see Fig. S7), resulting in a F_v_/F_m_ dynamics that follows the response of the approximately simulated exponential decay of PSII more closely. Thus, our theoretical analysis allowed discrimination between the effects of higher energy quenching of inactive PSII and energy transfer. Our results support the existence of energy transfer processes from active to inactive PSII.

In conclusion, we used a mathematical model of the PETC to investigate the fluorescence signal during photoinhibition and identified key factors that need to be included in order to realistically explain experimental fluorescence data. In addition to the hypotheses explored in this work, there are many other conceivable extensions and improvements. One possible extension is to include PSI fluorescence, as was done in [38]. We speculate that the PSI contribution might lead to a more realistic reproduction of the F_o_ signal. In addition, it may become important to include a description of PSII heterogeneity. The PSII pool consists of so called PSII*α* and PSII*β* complexes. Both differ in their antenna size and localization in the thylakoid membrane [24, 4]. In preliminary investigations we found that including such a heterogeneity does not change the slope ratio as defined in Eq. (13), which is a key indicator for the model response (see supplement). However, a full and realistic implementation of PSII*α* and PSII*β* and their different properties into our dynamic model is a future project. So far, also spatial effects have been ignored, in order to reduce the complexity of the *in silico* analysis. However, considering the complex three-dimensional structure of thylakoid membranes, these may be important to consider for more realistic models [13]. Additionally, it has been shown that the spatial architecture of leaves and the place of measurement (ad-, abaxial, or within leaves) influence the fluorescence signal obtained by spectroscopic techniques during photoinhibition [30]. Because we used a Dual-KLAS-NIR device for our measurements that records fluorescence on the abaxial leaf surface, future model versions should account for different local origins of the fluorescence signal. This is because the changes in the fluorescence signal obtained by devices measuring the abaxial surface, such as a Dual-KLAS-NIR, might correlate more with changes in chloroplasts in the lower than in the upper layers of the leaf. We envisage that our model can be used as a platform for the investigation of photoinhbitory effects, with several applications in mind. These include the study of long-term extinction phenomena (qZ and qH), which could support experimental efforts to identify the molecular mechanisms responsible for such quenching phenomena [20]. Moreover, our model also opens the possibility of investigating evolutionary questions. For example, by modifying the appropriate parameters, it can be used to explore the quenching capacities of a wide range of plant and algal species, thus supporting the generation of hypotheses explaining the enormous natural variation found in photoprotective processes [22, 35].

## Methods

A mathematical model was developed that combines non-photochemical quenching, the D1 protein repair cycle, and the main protein complexes in the PETC. The model is based on published mathematical descriptions that successfully simulated experimental data in the past [40, 6, 22]. Most parameter values were obtained from the literature. The model was tested against published data from various plant species and experimentally measured F_v_*/*F_m_ values (*Arabidopsis thaliana* ecotype Columbia-0 and the *npq1* mutant).

### Experimental approach

*Arabidopsis thaliana* (Columbia-0 and *npq1*) seeds were sown on commercial soil (Pikier, Balster Einheitserdewerk, Fröndenberg, Germany) and stratified for three days in the dark at 4 ^*°*^C. After that, they were transferred to the climate chamber with 12 h*/*12 h light/dark photoperiod, 26 ^*°*^C*/*20 ^*°*^C day/night air temperature and 60% relative air humidity. The photosynthetically active radiation was provided by fluorescent lamps (Fluora L58 W/77; Osram, Munich, Germany) with an intensity of approximately 100 µmol m^−2^ s^−1^at plant height. Finally, seedlings were transferred to pots (7*×* 7 *×*8 cm, one plant per pot) filled with soil (Lignostrat Dachgarten extensive, HAWITA, Vechta, Germany). Care was taken to avoid soil drying during cultivation. Six to seven weeks old plants were used for measuring.

Leaves of *A*.*thaliana* plants were detached, and petioles were submerged in a 5 mM lincomycin solution in reaction tubes for 3 h in dim light under ventilation. After incubation in the lincomycin solution, leaf discs with a diameter of 1.1 cm were punched out and floated on a water bath to keep the leaf temperature constant at 20 ^*°*^C. The floating leaf discs were exposed to white LED light (SL 3500-W-G, Photon Systems Instruments) with an intensity of 800 µmol m^−2^ s^−1^. After 0 h, 0.5 h, 1 h, 3 h, 5 h, and 6 h hours, F_v_/F_m_ was measured on six replicate leaf discs using a DUAL-KLAS-NIR system (Heinz Walz GmbH, Effeltrich, Germany). Each leaf was dark-adapted 20 minutes before a red saturation pulse (635 nm, 0.8 seconds) of >10000 µmol m^−2^ s^−1^was applied from both upper and lower sides of the leaf. Fluorescence was detected on the lower leaf surface to determine F_m_.

### Model description

Simulations were based on previous models of photosynthesis [6, 22] and the D1 protein repair cycle. For a detailed explanation, see the supplement. The photosynthetic electron transport chain in the thylakoid membrane of chloroplasts is implemented according to [6]. A four-state Photosystem II (PSII) description (*B*_0_ open and non-excited, *B*_1_ open and excited, *B*_2_ closed and non-excited, *B*_3_ closed and excited) was used. The rate of cytochrome b_6_f complex is described via mass-action kinetics. Photosystem I (PSI) is a three-state system similar to PSII. Convenience kinetics describes the activities of the ferredoxin-NADPHreductase (FNR) [19]. The proton leak across the thylakoid membrane, ATP synthesis, and cyclic electron flow around PSI are modeled via mass action kinetics. Reversible reactions are included by calculating lumenal pH-dependent equilibrium constants. Similar to [23] and [36], a four-state quencher module, based on the xanthophyll cycle and the protonation of PsbS, was integrated (see Fig. 1). The model is detailed in the supplementary material.

### D1 protein repair cycle and fluorescence

The repair and synthesis of the D1 protein of PSII were implemented by first-order equations governing the dynamics of three states of PSII [40]. These are PSII with intact D1 protein (U_a_), PSII with damaged D1 protein (U_i_), and PSII without D1 protein (U_d_). Here *U*_*a*_ = ∑_*i* …=1 4_ *B*_*i*_ comprises the four states of the model without photoinhibition.

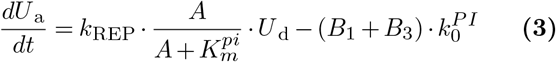

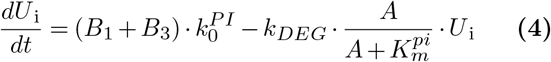

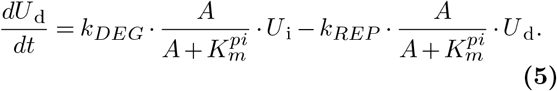

Here k_REP_ and k_DEG_ are the rate constants for the insertion of newly synthesized and degradation of damaged D1 protein. k^PI^ is the rate constant of photoinhibition. Several studies indicate that photoinhibition is a costly, energy-consuming process [34, 27]. Hence, degradation and insertion (PSII repair) of the D1 protein is proportional to the ATP concentration.

### Fluorescence

We assume that inactive PSII can dissipate excitation energy as heat and emit fluorescence. The fluorescence emitted by these PSII states is still affected by quenching.

#### Isolated PSII

Assuming no energy transfer between active and inactive PSII, the yield of fluorescence is described as (see [7, 6]),

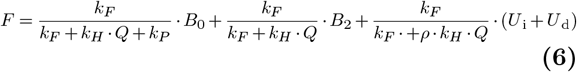

Here k_F_, k_P_, and k_H_ are the rate constant of fluorescence, photochemistry, and dissipitation of light energy other than fluorescence and photochemistry. *B*_0_ and *B*_2_ are open and closed states of active PSII (U_a_). The parameter *ρ* has been introduced to account for different heat dissipation properties between active and inactive PSII. Specifically, it describes the ratio of energy dissipation rates as heat between inactive (*U*_*i*_ + *U*_*d*_) and active (*U*_*a*_ the quencher activity.

Minimal fluorescence (*F*_*o*_) is observed in a dark-adapted state, where *B*_0_ *≈ U*_*a*._ Thus,

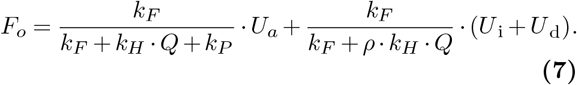

Assuming there are no inactive photosystems, Eq. (7) becomes,

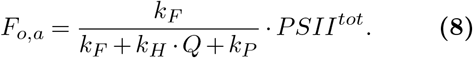

This is the expected *F*_*o*_ signal at the beginning of an experiment before high-light treatment started.

The maximal fluorescence yield is obtained in saturating light conditions, where *B*_2_ *≈U*_*a*_. Therefore,

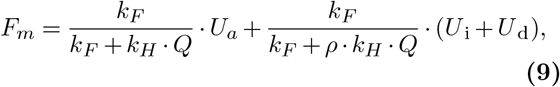

and without inactive PSII, representing the signal at the beginning of high-light treatment,

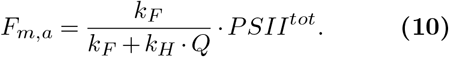

To quantify the response of *F*_*o*_ and *F*_*m*_ stress, we determine the derivatives of the relative fluorescence signals with respect to the active reaction centres, *U*_*a*_. The non-inhibited state corresponds to = PSII^tot^. We define

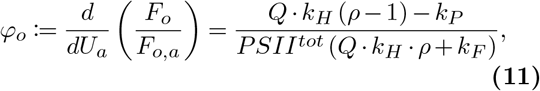

and

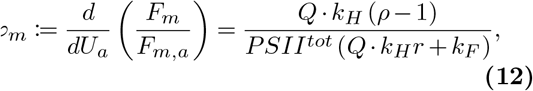

the ratio of these two values,

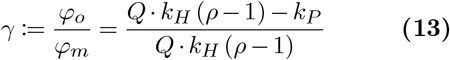

For a non-photoinhibited state, we get with Eqs. (8) and (10)

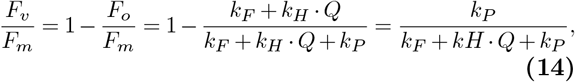

and, likewise using Eqs. (7) and (9), for a photoinhibited state

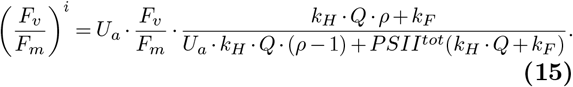

Eq. 15 becomes Eq. 14 when U_a_ = *PSII*^*tot*^.

#### Connected inactive and active PSII

In a second model, we assume that active closed PSII can transfer excitation energy to damaged PSII, see [7]. We describe this energy transfer rate as a first order process with rate constant *k*_*T*_ . This leads to the following description of the fluorescence signal,

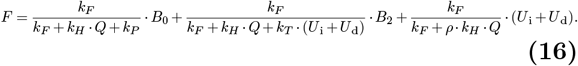

Hence,

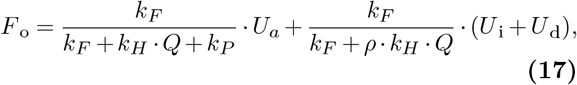

and

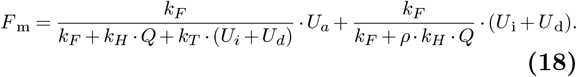

The expression for F_m_ is a rational function of active PSII (*U*_*i*_ + *U*_*d*_ = *PSII*^*tot*^*− U*_*a*_). This function has a singularity at,

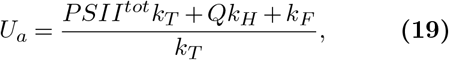

and extrema at,

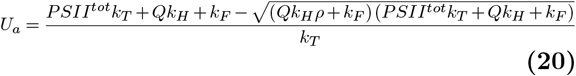

as well as,

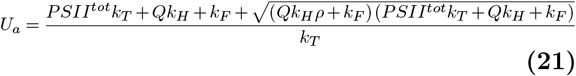

Note that for *k*_*T*_ =0 the expressions for F_m_ and F_o_ are identical to the isolated case. Using Eqs. 17 and 18 we can derive an expression for F_v_/F_m_,

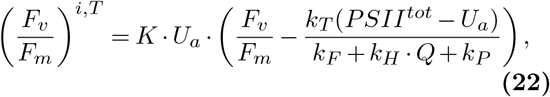

where *K*

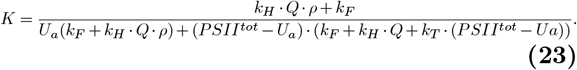

For *k*_*T*_ =0 Eq. 22 becomes identical to Eq. 15.

### ATP source

In previous models [6, 23], an external influx of ATP into the chloroplast is not included. However, several studies have shown that the metabolism of chloroplasts and mitochondria are interconnected and can influence each other [8, 42, 43]. We assumed that during light conditions, the external influx of ATP into the chloroplast is negligible, and the activity of the PETC provides all ATP. We model the external influx of ATP as constant flux with a light switch to ensure the resynthesis of the D1 protein in darkness.

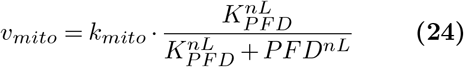

### Computational analysis

The model was implemented in the Python-based software modelbase version 1.3.8 [41]. For simulations the cvode solver implemented in Assimulo [1] was used. Python files containing the model and analyses can be found in the Gitlab repository https://gitlab.com/qtb-hhu/models/2023-photoinhibition.

## Abbreviations

CBB: Calvin-Benson-Bassham-cycle
PETC: photosynthetic electron transport chain
ROS: reactive oxygen species
U_a_: active photosystem II
U_i_: damaged photosystem II
U_d_: D1 protein-less photosystem II
PSII: photosystem II.

## Acknowledgment

We thank Ana Carolina dos Santos Sá and Yuxi Niu for their help during the experimental measurements.

## Author contributions

TN, OE: initial idea and conceptualisation. OE: funding acquisition. TN: visualisation. TN: formal analyses. TN, OE: writing—original draft and introduction. TN, OE: writing—original draft and methods. TN,OE: writing—original draft and results. TN, OE: writing—original draft, discussion, and TN, OE, SM writing—review and editing. All authors read and accepted the final version of the manuscript.

## Data Availability Statement

The original contributions presented in the study are included in the article/Supplementary Material, further inquiries can be directed to the corresponding author/s. The code can be found at https://gitlab.com/qtb-hhu/models/2023photoinhibition

