## Supplemental Text for "A mathematical model of photoinhibition: exploring the impact of quenching processes"

### Model

The mathematical model of the photosynthetic electron transport chain, including non-photochemical quenching and the D1 protein repair cycle, is represented as a system of twelve ordinary differential equations. The equations describe the temporal evolution of twelve system variables. These variables are the reduced fraction of plastoquinone (P), plastocyanin (PC), and ferredoxin (Fd), the stromal concentration of ATP (A) and NADPH (N), the lumenal proton concentration (H), the oxidized fraction of PsbS and the fraction of Violaxanthin, the active ($U_a$), damaged ($U_i$), and D1 protein less ($U_d$) form of photosystem II as well as an activator of the ATP synthase (E). Most of the equations are taken from previous work [1–3]. The system of equations reads,

$$\frac{dPQH_2}{dt} = v_{PSII} + v_{cyc} - v_{b6f} - v_{PTOX} \tag{1}$$

$$\frac{dPC^-}{dt} = 2\ v_{b6f} - v_{PSI} \tag{2}$$

$$\frac{dFd^-}{dt} = v_{PSI} - 2\ v_{FNR} - 2\ v_{cyc} \tag{3}$$

$$\frac{dN}{dt} = v_{FNR} - v_{NADPHconsumption} \tag{4}$$

$$\frac{dA}{dt} = v_{ATPsynthase} + v_{mito} - v_{ATPconsumption} - 231\ v_{deg} - 1059\ v_{rep} \tag{5}$$

$$b_H \cdot \frac{dH}{dt} = 2\ v_{PSII} + 4\ v_{b6f} - \frac{14}{3}\ v_{ATPsynthase} - v_{leak} \tag{6}$$

$$\frac{dPsbS}{dt} = v_{Deprot} - v_{Prot} \tag{7}$$

$$\frac{dV}{dt} = v_{EpoxZ} - v_{DeepoxV} \tag{8}$$

$$\frac{dU_a}{dt} = v_{rep} - v_{inh} \tag{9}$$

$$\frac{dU_i}{dt} = v_{inh} - v_{deg} \tag{10}$$

$$\frac{dU_d}{dt} = v_{deg} - v_{rep} \tag{11}$$

$$\frac{dE}{dt} = v_{actATPsyn} - v_{deactATPsyn} \tag{12}$$

#### Photosystems and Fluorescence

As outlined in [2], we assume processes in photosystem II (PSII) and photosystem I (PSI) to be much faster than the rest of the photosynthetic electron transport chain. This assumption allows us to treat the photosynthetic electron transport chain as a fast-slow system and algebraically solve the equations governing the dynamics in the photosystems. Hence, Photosystem II is represented as a four-state, and PSI is formulated as a three-state algebraic equation system. The equation system for PSII, in the case of no energy transfer between inactive and active PSII, is

$$-\left(k_{LII}+\frac{k_{PQred}}{K_{eq,QAPQ}}PQH_2\right)B_0+(k_H+k_F)B_1+k_{PQred}PQ\cdot B_2=0 \tag{13}$$

$$k_{LII}B_0-(k_H+k_F+k_P)B_1=0 \tag{14}$$

$$k_{LII}B_2-(k_H+k_F)B_3=0 \tag{15}$$

$$B_0+B_1+B_2+B_3=U_a. \tag{16}$$

Thus, $v_{PSII}$ is (see [4])

$$v_{PSII}=0.5\cdot k_P\cdot B_1. \tag{17}$$

Here $\mathrm{k_P}$, $\mathrm{k_F}$ and $\mathrm{k_H}$ are the rate constants of photochemistry, fluorescence, and heat dissipation, respectively. The parameter $\mathrm{k_{PQred}}$ describes plastoquinone reduction at Photosystem II.

Four states of active Photosystem II ($\mathrm{U_{act}}$) are considered in the solution of the PSII equation system. Two ground and two excited states of open ($\mathrm{B_0}$, $\mathrm{B_1}$) and closed ($\mathrm{B_2}$, $\mathrm{B_3}$) reaction centers. The states differ in the ways they use light energy.

With the assumption that inactive PSII can emit excitation energy as fluorescence and heat (with higher or lower efficiency than active PSII), the fluorescence signal calculates as,

$$F=\frac{k_F}{k_F+k_H\cdot Q+k_P}\cdot B_0+\frac{k_F}{k_F+k_H\cdot Q}\cdot B_2+\frac{k_F}{k_F+\rho\cdot k_H\cdot Q}\cdot(U_i+U_d). \tag{18}$$

Note that we approximate open and closed PSII with $B_0$ and $B_2$, because $B1$ and $B3$ acquire only small values during fluorescence transients.

The equations governing the dynamics in a system with energy transfer between active and inactive PSII reads,

$$-\left(k_{LII}+\frac{k_{PQred}}{K_{eq,QAPQ}}PQH_2\right)B_0+(k_H\cdot Q+k_F)B_1+k_{PQred}PQ\cdot B_2=0 \tag{19}$$

$$k_{LII}B_0-(k_H\cdot Q+k_F+k_P)B_1=0 \tag{20}$$

$$(k_H\cdot Q+k_F)B_3+k_PB_1+\frac{k_{PQred}}{K_{eq,QAPQ}}PQH_2B_0-(k_{PQred}PQ+k_{LII})B_2+k_T(U_{in}+U_{dn})B_3=0 \tag{21}$$

$$B_0+B_1+B_2+B_3=U_a \tag{22}$$

$$k_{LII}U_{in}-(\rho\cdot k_H\cdot Q+k_F)U_{ip}+k_TB_3U_{in}=0 \tag{23}$$

$$U_{ip}+U_{in}=U_i \tag{24}$$

$$k_{LII}U_{dn}-(\rho\cdot k_H\cdot Q+k_F)U_{dp}+k_TB_3U_{dn}=0 \tag{25}$$

$$U_{dp}+U_{dn}=U_d \tag{26}$$

$$\tag{27}$$

and the fluorescence signal calculates as,

$$F=\frac{k_F}{k_F+k_H\cdot Q+k_P}\cdot B_0+\frac{k_F}{k_F+k_H\cdot Q+k_T(U_i+U_d)}\cdot B_2+\frac{k_F}{k_F+\rho\cdot k_H\cdot Q}\cdot(U_i+U_d). \tag{28}$$

The equation system for PSI is,

$$0=\frac{dY_0}{dt}=k_{PCox}\cdot PC^-\cdot Y_2-\frac{k_{PCox}}{K_{eq,PCP700}}\cdot PC\cdot Y_0-k_{LI}Y_0 \tag{29}$$

$$0=\frac{dY1}{dt}=k_{LI}Y_0-k_{Fdred}\cdot Fd\cdot Y_1+\frac{k_{PCox}}{K_{eq,P700Fd}}\cdot Fd^-\cdot Y_2 \tag{30}$$

Assuming that the total amount of PSI is conserved,

$$PSI^{tot} = Y_0 + Y_1 + Y_2, \tag{31}$$

The equation system for PSI can be solved to obtain steady state expressions of the PSI states.

The rate of PSI is,

$$v_{PSI} = k_{LI} \cdot Y_0. \tag{32}$$

### Cytochrome $b_6$f

The reaction mechanism of Cytochrome $b_6$f is formulated as simple mass-action kinetics:

$$v_{b6f} = max\left(k_{b6f} \cdot \left(PQH_2 \cdot PC^2 - \frac{PQ \cdot PC^-}{K_{eq,b6f(H)}}\right), v_{b6f}^{min}\right) \tag{33}$$

### Ferredoxin-NADPH reductase

The Ferredoxin-NADPH is described as convenience kinetics (see [2, 5]). s:

$$v_{FNR} = V_{FNR}^{max} \cdot \frac{f^{-2} \cdot n^+ - (f^2 \cdot n)/K_{eq,FNR}}{(1 + f^- + f^{-2}) \cdot (1 + n^+) + (1 + f + f^2) \cdot (1 + n^+) - 1} \tag{34}$$

here, $f$, $f^-$,$n^+$ and n are:

$$f = \frac{Fd}{K_{M,F}},\ f^- = \frac{Fd^-}{K_{M,F}},\ n^+ = \frac{NADP^+}{K_{M,N}},\ n = \frac{NADPH}{K_{M,N}} \tag{35}$$

### Cyclic electron flow

The model includes a simplified description of cyclic electron transport around photosystem I.

$$v_{cyc} = k_{cyc} \cdot (Fd^-)^2 \cdot PQ, \tag{36}$$

Following [2] we assume the cyclic electron flow via the reduction of the plastoquinone pool by ferredoxin to be irreversible and for various cyclic electron flow reactions one combined reaction.

### ATPsynthase

For the rate of ATP synthase, the description presented in [2, 6] was used,

$$v_{ATPsyn} = k_{ATPsyn} \cdot E \cdot \left(ADP - \frac{ATP}{K_{eq,ATPsyn}(H)}\right) \tag{37}$$

.

A pH-dependent activation ATP synthase was also included (compare [3]). The activation rate is

$$v_{actATPsyn} = k_{actATPsyn} \cdot H(PFD) \cdot E. \tag{38}$$

Deactivation is formulated as

$$v_{deactATPsyn} = k_{deactATPsyn} \cdot (1 - H(PFD)) \cdot (1 - E). \tag{39}$$

Here H(PFD) is a function that is zero in dark and one in light (PFD $\geq 1$) conditions.

### D1 protein repair cycle

A simple D1 protein repair cycle model was used for simulating photoinhibition effects [1]. The equations describing the state changes between active, damaged, and D1 protein-less PSII are

$$\frac{dU_a}{dt} = k_{REP} \cdot \frac{A}{A + K_m^{pi}} \cdot U_d - (B1 + B3) \cdot k_0^{PI} \tag{40}$$

$$\frac{dU_i}{dt} = (B1 + B3) \cdot k_0^{PI} - k_{DEG} \cdot \frac{A}{A + K_m^{pi}} \cdot U_i \tag{41}$$

$$\frac{dU_d}{dt} = k_{DEG} \cdot \frac{A}{A + K_m^{pi}} \cdot U_i - k_{REP} \cdot \frac{A}{A + K_m^{pi}} \cdot U_d. \tag{42}$$

The ATP-dependency of the degradation and repair processes [7] are included via a Hill-like activation term.

### Xanthophyll cycle

The representation of the xanthophyll cycle follows [3]. This description of the xanthophyll cycle only considers Violaxanthin and Zeaxanthin while ignoring Antheraxanthin.

The deepoxidation from Violaxanthin to Zeaxanthin is,

$$v_{VDP} = k_{deepox} \cdot \frac{H^{n_{HX}}}{H^{n_{HX}} + pH_{inv}(K_{sat})^{n_{HX}}} \cdot V. \tag{43}$$

The epoxidation rate from Zeaxanthin to Violaxanthin is,

$$v_{ZEP} = k_{epox} \cdot Z \tag{44}$$

### PsbS protonation and deprotonation

Equations of protonation and deprotonation of PsbS are used from [3],

$$v_{prot} = k_{prot} \cdot \frac{H^{n_{HL}}}{H^{n_{HL}} + pH_{inv}(K_{sat,PsbS})^{n_{HL}}} PsbS \tag{45}$$

and

$$v_{deprot} = k_{deprot} PsbS^p. \tag{46}$$

### Quencher

Quenching is formulated via a four-state quencher module. This module includes the Xanthophyll cycle and the protonation state of PsbS. Each state is weighted by a parameter ($\gamma_0$, $\gamma_1$, $\gamma_2$, $\gamma_3$) and the Quencher activity is then calculated as,

$$Q = \gamma_0 \cdot (1 - Z_s) \cdot PsbS + \gamma_1 \cdot (1 - Z_s) \cdot PsbS^p + \gamma_2 \cdot Z_s \cdot PsbS^p + \gamma_3 \cdot Z_s \cdot PsbS, \tag{47}$$

where the contribution of Zeaxanthin ($Z_s$) is (compare [3]),

$$Zs = \frac{Z}{Z + Kzsat} \tag{48}$$

Additionally, when considering a long-term quenching compound, the simplified assumption was made that this compound's activity is proportional to the fraction of inactive PSII, resulting in a fifth term for the quencher's activity.

$$\gamma_4 \cdot \left(1 - \frac{U_a}{PSII^{tot}}\right). \tag{49}$$

### ATP and NADPH consumption

All processes that consume ATP and NADPH (except the resynthesis of the D1 protein) are combined into one term each,

$$v_{ATPcons} = k_{ATPcons} \cdot ATP \tag{50}$$

and

$$v_{NADPHcons} = k_{NADPHcons} \cdot NADPH \tag{51}$$

An external ATP flux must be assumed to ensure repair of damaged PSII prolonged dark conditions

$$v_{mito} = k_{mito} \cdot \frac{K_{PFD}^{nL}}{K_{PFD}^{nL} + PFD^{nL}} \tag{52}$$

### Proton leak

The thylakoid membrane is assumed to be leaky for protons [2, 3].

$$v_{leak} = k_{leak} \cdot (pH - pH_{stroma}) \tag{53}$$

### PTOX

For the implementation of PTOX a constant stromal oxygen concentration is assumed ($O_2^{ext}$).

$$v_{PTOX} = k_{PTOX} \cdot O_2^{ext} \cdot PQ \tag{54}$$

### Equilibrium constants

To include the electrical and pH-dependent contribution of the protonmotive force on the activities of the PETC, we calculated the equilibrium constants as outlined initially in [2] and used in [3, 4, 8]. We repeat the example as outlined in [2] for clarification. the overall reaction for cytchrome $b_6f$ is

$$PQH_2 + 2\,PC + 2\,H^+_{stroma} \rightleftharpoons PQ + 2\,PC^- + 4\,H^+_{lumen}.$$

We split the reaction into two redox half-reactions and one transport process:

$$\text{(a)}\ PQ + 2\,e^- + 2\,H^+_{lumen} \rightleftharpoons PQH_2$$

$$\text{(b)}\ PC + e^- \rightleftharpoons PC^-$$

$$\text{(c)}\ H^+_{stroma} \rightleftharpoons H^+_{lumen}$$

The overall reaction is the stoichiometric sum

$$-1 \cdot (a) + 2 \cdot (b) + 2 \cdot (c). \tag{55}$$

The contribution to the standard Gibbs free energy change are

$$\Delta G_1^o = -2FE_o(PQ/PQH_2) + 2RT\ ln(10) \cdot pH_{lumen} \tag{56}$$

$$\Delta G_2^o = -2FE_0(PC/PC^-) \tag{57}$$

$$\Delta G_3^o = RT\ ln(10)(pH_{stroma} - pH_{lumen}). \tag{58}$$

Hence, according to the stoichiometric sum the overall standard Gibbs free energy change amounts to

$$\Delta G^o = -\Delta G_1^o + 2\Delta G_2^o + 2\Delta G_3^o \tag{59}$$

The equilibrium constant can be easily derived using $\Delta G^o$.

### Additional figures

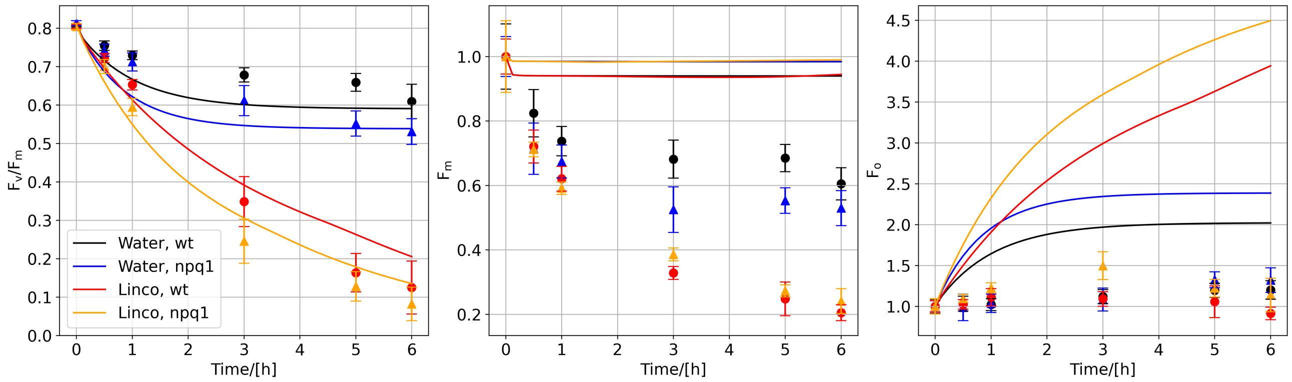


Figure S1: Experimental measurement and simulated changes in $F_v/F_m$, $F_m$, and $F_o$ in high light treatment of *A.thaliana* plants for 6 hours. The plants were treated with water (black and blue lines) or lincomycin (red and orange lines), inhibiting protein synthesis. Light intensity was 800 $\mu mol\, m^{-2}\, s^{-1}$. The model did not include energy transfer processes and difference in heat dissipation properties between active and inactive PSII.

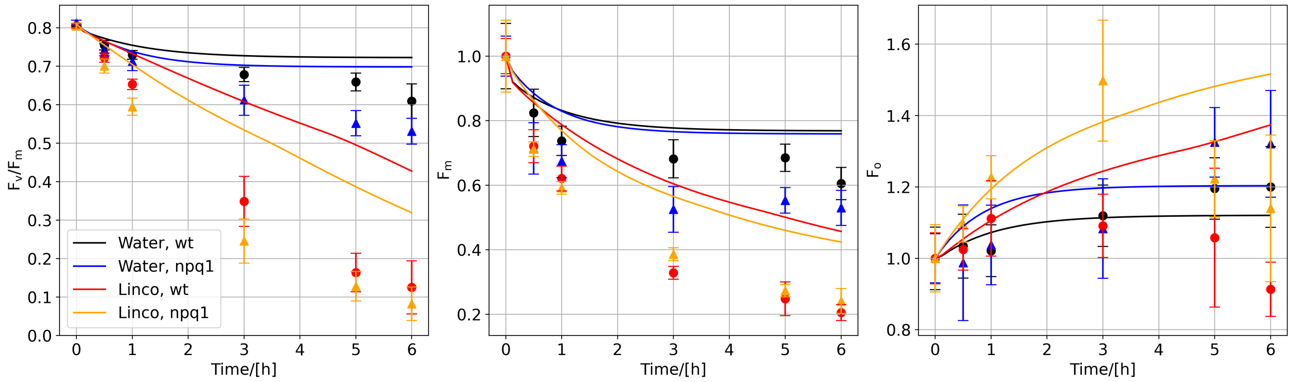


Figure S2: Experimental measurement and simulated changes in $F_v/F_m$, $F_m$, and $F_o$ in high light treatment of *A.thaliana* plants for 6 hours. The plants were treated with water (black and blue lines) or lincomycin (red and orange lines), inhibiting protein synthesis. Light intensity was 800 $\mu mol\, m^{-2}\, s^{-1}$. The model did include differences in heat dissipation properties between active and inactive PSII ($\rho$=6).

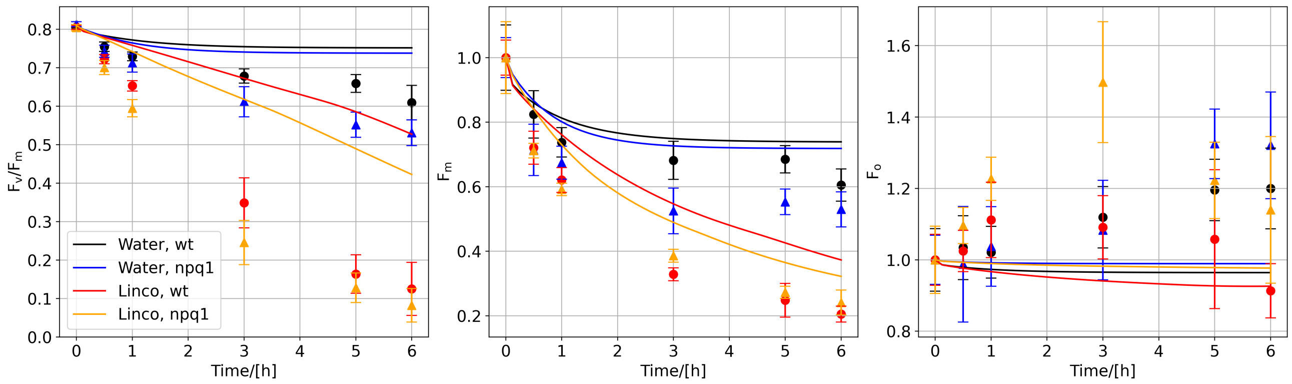


Figure S3: Experimental measurement and simulated changes in $F_v/F_m$, $F_m$, and $F_o$ in high light treatment of *A.thaliana* plants for 6 hours. The plants were treated with water (black and blue lines) or lincomycin (red and orange lines), inhibiting protein synthesis. Light intensity was 800 µmol $m^{-2}$ $s^{-1}$. The model did include differences in heat dissipation properties between active and inactive PSII ($\rho$=11).

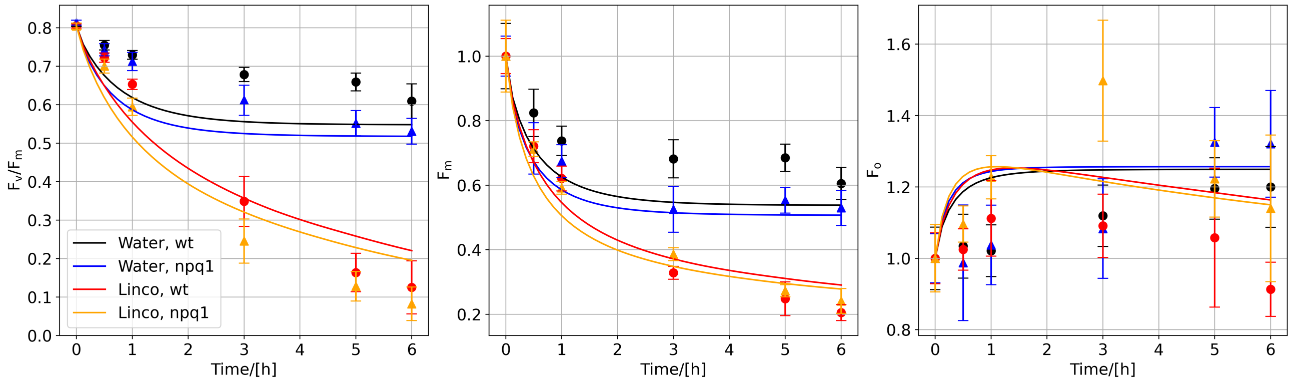


Figure S4: Experimental measurement and simulated changes in $F_v/F_m$, $F_m$, and $F_o$ in high light treatment of *A.thaliana* plants for 6 hours. The plants were either treated with water (black and blue lines) or lincomycin (red and orange line) inhibiting protein synthesis. Light intensity was 800 µmol $m^{-2}$ $s^{-1}$. $\rho = 1$, $kT = 0$, and slow quenching was used.

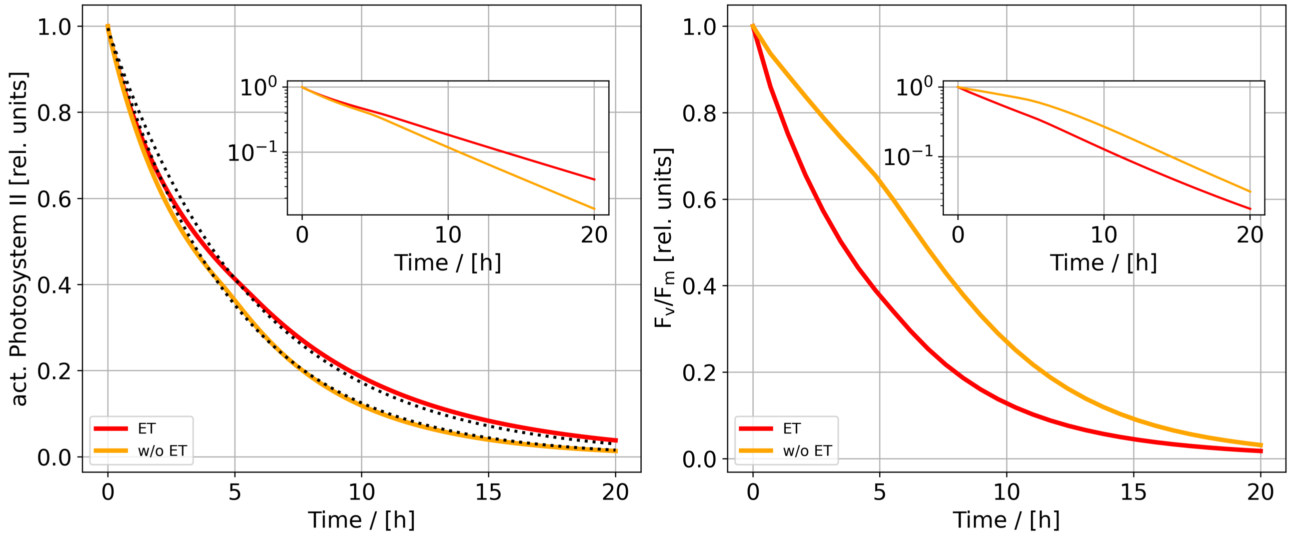


Figure S5: Simulated changes of active PSII and $F_v/F_m$ in a model version with (red) and without (orange) energy transfer between active and inactive PSII ($k_T$=0) during high light treatment (800 µmol $m^{-2}$ $s^{-1}$). The simulation were supposed to happen in lincomycin-treated tissue. Dotted lines indicate a fit to exponential function. Insets show the same data but on logarithmic scale.

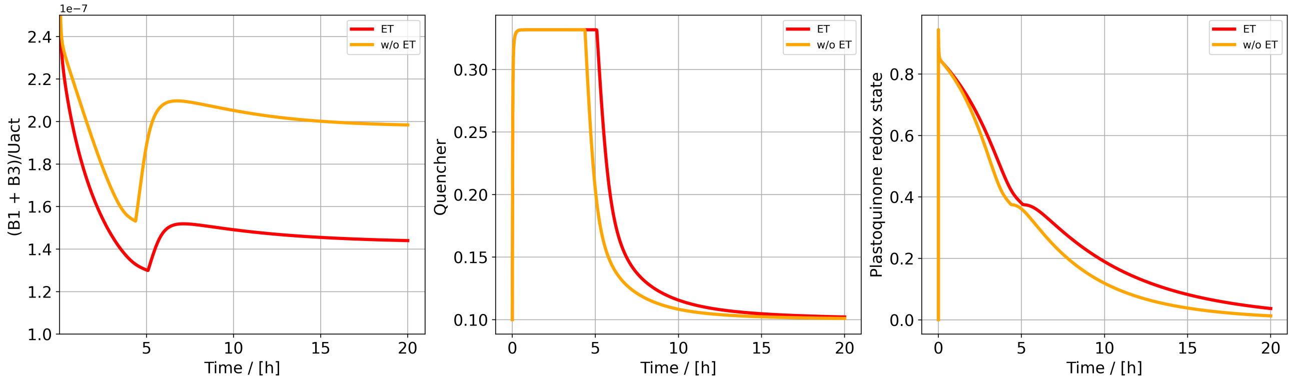


Figure S6: Simulated changes of excited active PSII states (B1 and B3), quencher activity and plastoquinone redox state in a model version with (red) and without (orange) energy transfer between active and inactive PSII ($k_T$=0) during high light treatment (800 µmol m$^{-2}$ s$^{-1}$). The simulation were supposed to happen in lincomycin-treated tissue.

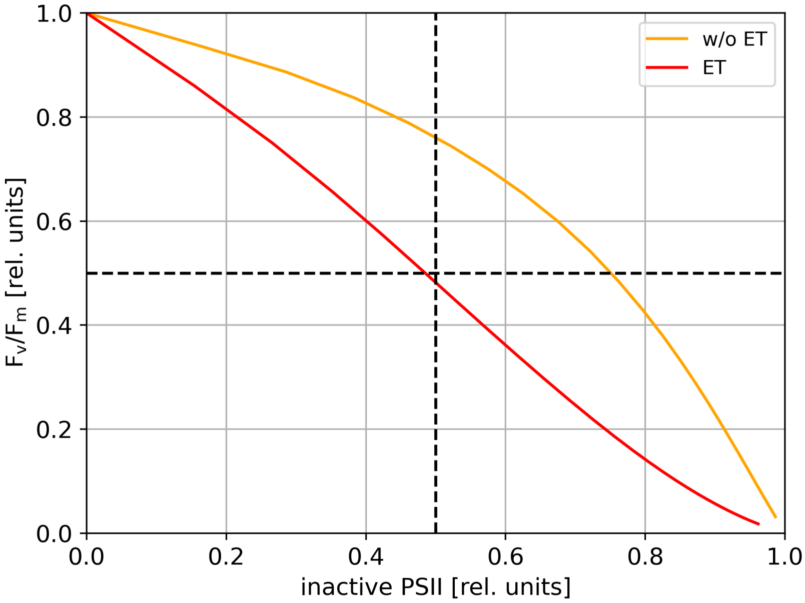


Figure S7: Simulated changes of $F_v/F_m$ as function of inactive PSII in a model version with (red) and without (orange) energy transfer between active and inactive PSII ($k_T$=0) during high light treatment (800 µmol m$^{-2}$ s$^{-1}$). The simulation were supposed to happen in lincomycin-treated tissue.

### Connected units with forth- and back transfer of energy between active and damaged PSII

For our theoretical approach, we assume that energy transfer can happen between closed and open reaction centers and between damaged and active reaction center no matter in which state they are. For simplicity we assume that energy transfer does not happen for active open reaction centers and the energy transfer between the active closed reaction centers and damaged reaction centers is the same in both directions. This leads to following description of the fluorescence signal,

$$F = \frac{k_F}{k_F + k_H \cdot Q + k_P} \cdot B_0 + \frac{k_F}{k_F + k_H \cdot Q + kT \cdot B_0 + k_{T2} \cdot (U_\mathrm{i} + U_\mathrm{d})} \cdot B_2 + \quad (60)$$

$$\frac{k_F}{k_F \cdot +\rho \cdot k_H \cdot Q + k_{T2} \cdot (B_0 + B_2)} \cdot (U_\mathrm{i} + U_\mathrm{d}). \quad (61)$$

Hence, $F_o$ is,

$$F_o = \frac{k_F}{k_F + k_H \cdot Q + k_P} \cdot U_a + \frac{k_F}{k_F \cdot + \rho \cdot k_H \cdot Q + k_{T2} \cdot U_a} \cdot (U_i + U_d), \tag{62}$$

and $F_m$,

$$F_m = \frac{k_F}{k_F + k_H \cdot Q + k_{T2} \cdot (U_i + U_d)} \cdot U_a + \frac{k_F}{k_F \cdot + \rho \cdot k_H \cdot Q + k_{T2} \cdot U_a} \cdot (U_i + U_d). \tag{63}$$

Using equation 62 and 63 we can derive an expression for $F_v/F_m$,

$$\left(\frac{F_v}{F_m}\right)^{i,T} = K \cdot U_a \cdot \left(\frac{F_v}{F_m} - \frac{k_{T2}(PSII - U_a)}{k_F + k_H \cdot Q + k_P}\right), \tag{64}$$

where $K$

$$K = \frac{k_H \cdot Q \cdot \rho + U_a \cdot k_{T2} + k_F}{U_a(k_F + k_H \cdot Q \cdot \rho + k_{T2} \cdot U_a) + (PSII - U_a) \cdot (k_F + k_H \cdot Q + k_{T2} \cdot (PSII - Ua))}. \tag{65}$$

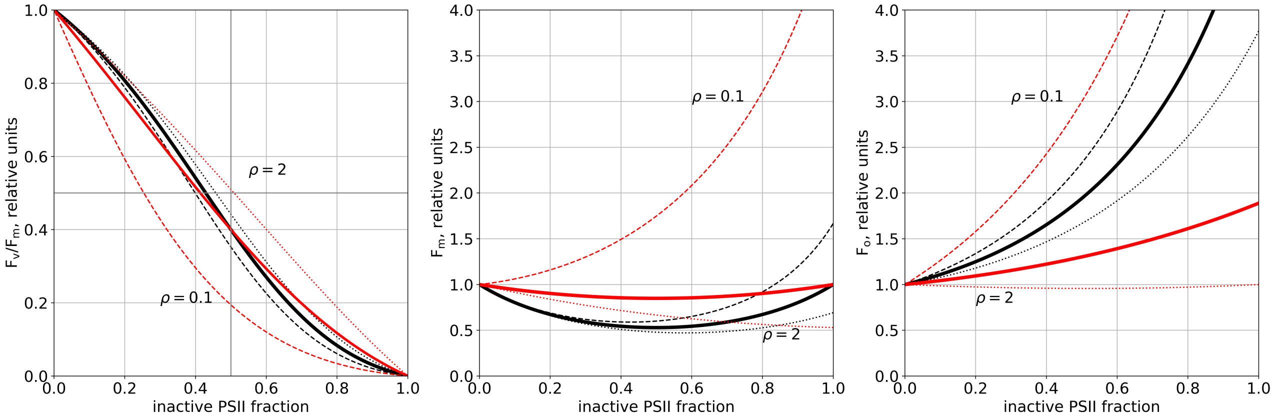


Figure S8: Relationship between $F_v/F_m$, $F_m$, and $F_o$ and fraction of active photosystem II in a model with energy transfer. The dashed, continuous, and dotted lines indicate scenarios in which the ratio of heat dissipation between active and inactive PSII is 0.1, 1, and 2, respectively. Black lines signify a low quenching, while red lines denote a high quenching activity (Q = 0.1 and 1). Parameter values used for the calculations can be found in the parameter table.

### PSII heterogeneity

According to [9], PSII heterogeneity could influence the fluorescence signal observed during photoinhibitory treatment. To check how the relation between $F_o$, $F_m$ and $F_v/F_m$ and inactive PSI changes when we assume a mixed population of PSII$\alpha$ and PSII$\beta$ we repeated our analysis in the main text (without energy transfer) implementing PSII heterogeneity according to [9].

Hence, $F_o$ is,

$$F_o^H = PSII^{tot}\,(1-a)\,\frac{k_F}{k_F + k_H \cdot Q + k_P} + a\left(\frac{k_F}{k_F + k_H \cdot Q + k_P} \cdot U_a + \frac{k_F}{k_F + \rho \cdot k_H \cdot Q} \cdot (U_i + U_d)\right), \tag{66}$$

and $F_m$,

$$F_m^H = PSII^{tot}\,(1-a)\,\frac{k_F}{k_F + k_H \cdot Q} + a\left(\frac{k_F}{k_F + k_H \cdot Q} \cdot U_a + \frac{k_F}{k_F + \rho \cdot k_H \cdot Q} \cdot (U_i + U_d)\right). \tag{67}$$

Using the fact that $U_i + U_d = PSII^{tot} - U_a$, we can derive a formula for $\mathrm{F_v/F_m}$

$$\left(\frac{F_v}{F_m}\right)^{i,H} = \left(a\ U_a + (1-a)\ PSII^{tot}\right) \cdot \frac{F_v}{F_m} \cdot K, \tag{68}$$

where $K$ is

$$\frac{k_H \cdot Q \cdot \rho + k_F}{PSII^{tot}\ (1-a)\ (\rho \cdot k_H \cdot Q + k_F) + a\ (U_a\ (\rho \cdot k_H \cdot Q + k_F) + (PSII^{tot} - U_a)\ (k_H \cdot Q + k_F))}. \tag{69}$$

The slopes of the relative minimal ($\mathrm{F_o}$) and maximal fluorescence ($\mathrm{F_m}$) regarding active PSII ( $\mathrm{U_a}$) are

$$\varphi_o^H := \frac{d}{dU_a}\left(\frac{F_o^H}{F_{o,a}}\right) = \frac{a\ (Q \cdot k_H\,(\rho-1) - k_P)}{PSII^{tot}\,(Q \cdot k_H \cdot \rho + k_F)}, \tag{70}$$

and

$$\varphi_m^H := \frac{d}{dU_a}\left(\frac{F_m^H}{F_{m,a}}\right) = \frac{a \cdot Q \cdot k_H\,(\rho-1)}{PSII^{tot}\,(Q \cdot k_H r + k_F)}. \tag{71}$$

Hence, the slope ratio is

$$\gamma^H := \frac{\varphi_o^H}{\varphi_m^H} = \frac{Q \cdot k_H\,(\rho-1) - k_P}{Q \cdot k_H\,(\rho-1)}, \tag{72}$$

which is the same as in the isolated case (compare main text).

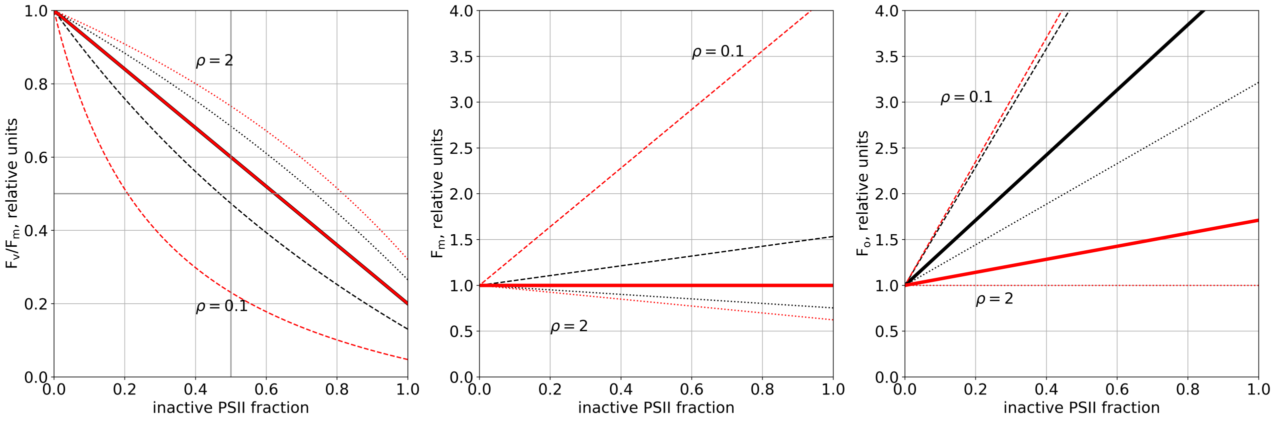


Figure S9: Relationship between $\mathrm{F_v/F_m}$, $\mathrm{F_m}$, and $\mathrm{F_o}$ and fraction of active photosystem II in a model with energy transfer and PSII heterogeneity. The dashed, continuous, and dotted lines indicate scenarios in which the ratio of heat dissipation between active and inactive PSII is 0.1, 1, and 2, respectively. Black lines signify a low quenching, while red lines denote a high quenching activity (Q = 0.1 and 1). Parameter values used for the calculations can be found in the table parameter table. a is set to 0.8.

### Parameters

Following table gives information about the parameters used and from which resource they were taken.

| parameter | value | reference/comment |
|---|---|---|
| Pool sizes | | |
| $PSII^{tot}$ | 2.5 mmol (mol Chl)$^{-1}$ | PSII reaction centers [10] |
| $PSI^{tot}$ | 2.5 mmol (mol Chl)$^{-1}$ | PSII reaction centers [10] |
| $PQ^{tot}$ | 20 mmol (mol Chl)$^{-1}$ | total plastoquinone pool [11] |
| $PC^{tot}$ | 4 mmol (mol Chl)$^{-1}$ | total plastocyanin pool [12] |
| $Fd^{tot}$ | 5 mmol (mol Chl)$^{-1}$ | total ferredoxin pool [12] |
| $ATP^{tot}$ | 50 mmol (mol Chl)$^{-1}$ | total adenosine phosphate pool [13] |
| $NADP^{tot}$ | 25 mmol (mol Chl)$^{-1}$ | $NADP^{+} + NADPH$ pool [13] |
| $PsbS^{tot}$ | 1 | normalized PsbS pool, after [3] |
| $X^{tot}$ | 1 | normalized total pool of xanthophylls, afetre [3] |
| Rate constants | | |
| $k_{ActATPase}$ | 0.01 s$^{-1}$ | activation of ATP synthase, after [3] |
| $k_{DeactATPase}$ | 0.002 s$^{-1}$ | deactivation of ATP synthase, after [3] |
| $k_{ATPsynthase}$ | 20 s$^{-1}$ | unchanged from [2] |
| $k_{ATPconsumption}$ | 10 s$^{-1}$ | unchanged from [2] |
| $k_{NADPHconsumption}$ | 20 s$^{-1}$ | after [2] |
| $k_H$ | $5 \cdot 10^9$ s$^{-1}$ | rate of non-radiative decay, after [3] |
| $k_F$ | $6.25 \cdot 10^8$ s$^{-1}$ | rate of fluorescence, after [3] |
| $k_P$ | $5 \cdot 10^9$ s$^{-1}$ | rate of photochemistry, after [3] |
| $k_{PTOX}$ | 0.01 mmol$^{-1}$ (mol Chl) s$^{-1}$ | after [2] |
| $k_{cyc}$ | 1 mmol$^{-2}$ (mol Chl)$^2$ s$^{-1}$ | after [2] |
| $k_{leak}$ | 100 s$^{-1}$ | |
| $k_{b6f}$ | 2.5 mmol$^{-2}$ (mol Chl)$^2$ s$^{-1}$ | after [2] |
| $v_{b6f}^{min}$ | -2.5 mmol (mol Chl)$^{-1}$s$^{-1}$ | after [2] |
| $k_{PCox}$ | 2500 mmol$^{-1}$ (mol Chl) s$^{-1}$ | after [2] |
| $k_{Fdred}$ | $2.5 \cdot 10^5$ mmol$^{-1}$ (mol Chl) s$^{-1}$ | after [2] |
| $k_{PQred}$ | 250 mmol$^{-1}$ (mol Chl) s$^{-1}$ | after [2] |
| $k_{PTOX}$ | 0.01mmol$^{-1}$ (mol Chl) s$^{-1}$ | after [2] |
| $V_{FNR}^{max}$ | 1500 mmol (mol Chl)$^{-1}$s$^{-1}$ | after [2] |
| $k_{Deepox}$ | 0.0024 s$^{-1}$ | rate of de-epoxidation, unchanged from [3] |
| $k_{Epox}$ | 0.00024 s$^{-1}$ | rate of epoxidation, unchanged from [3] |
| $k_{Deprot}$ | 0.0096 s$^{-1}$ | rate of PsbS de-protonation, unchanged from [3] |
| $k_{Prot}$ | 0.0096 s$^{-1}$ | rate of PsbS protonation, unchanged from [3] |
| $k_{DEG}$ | 0.0000833s$^{-1}$ | based on a degradation rate of 0.3/h [14] |
| $k_{PI0}$ | 300 s$^{-1}$ | ad-hoc estimation based on achieving comparable rates as when using the whole PSII pool and $k_{PI0}$=0.0044 $min^{-1}$ [1] |

| parameter | value | reference/comment |
|---|---|---|
| $k_{REP}$ | 0.00833 $s^{-1}$ | based on the observation that $U_d$ concentration is small in WT [14] |
| $K_{mPI}$ | 16 mmol (mol Chl)$^{-1}$ | |
| $k_T$ | $9 \cdot 10^8$ mmol$^{-1}$ (mol Chl) $s^{-1}$ | fitted to $F_v/F_m$ changes in high light |
| $\rho$ | 7 | fitted to $F_v/F_m$ changes in high light |
| $nL$ | 2 | ad-hoc value to use a reasonable switch behavior |
| $K_{PFD}$ | 100 | ad-hoc value to use a reasonable switch behavior |
| $mito$ | 100 mmol (mol Chl)$^{-1}$s$^{-1}$ | ad-hoc value, needs to be refined as model becomes more elaborate |
| Half-saturation and Michaelis constants | | |
| $K_{pHsatz}$ | 5.8 | half-saturation pH for de-epoxidase activity, after [3] |
| $K_{pHsatLHC}$ | 5.8 | pKa of PsbS activation, after [3] |
| $K_{Zsat}$ | 0.12 | after [3] |
| $K_{MF}$ | 1.56 mmol (mol Chl)$^{-1}$ | after [2] |
| $K_{MN}$ | 0.22 mmol (mol Chl)$^{-1}$ | after [2] |
| External concentrations | | |
| $O_2^{ex}$ | 8 mmol (mol Chl)$^{-1}$ | external oxygen pool after [2] |
| $P_i$ | 0.01 | internal pool of phosphates after [2] |
| Other constants | | |
| $F$ | 96.485 kJ | Farraday constant |
| $R$ | $8.3 \cdot 10^{-3}$ kJ $K^{-1}$ mol$^{-1}$ | universal gas constant |
| $T$ | 298 K | temperature |
| $\gamma_0$ | 0.1 | base quenching [3] |
| $\gamma_1$ | 0.25 | fast quenching due to protonation [3] |
| $\gamma_2$ | 0.6 | fastest possible quenching [3] |
| $\gamma_3$ | 0.105 | slow quenching by Zx [3] |
| $\gamma_4$ | 1.01 | test long-term quencher |
| $n_{HL}$ | 3 | Hill-coefficient for activity of de-protonation, after [3] |
| $n_{HX}$ | 5 | Hill-coefficient for deepoxidase activity, after [3] |
| $b_H$ | 100 | protonation buffering constant [15] |
| pHstroma | 7.8 | |
| Standard potentials | | |
| $E_0(Q_A/Q_A^-)$ | -0.140 V | unchanged from [2, 16] |
| $E_0(PQ/PQH_2)$ | 0.354 V | unchanged from [2, 17] |
| $E_0(PC/PC^-)$ | 0.380 V | unchanged from [2, 18] |

Table S1 – *Continued from previous page*

| parameter | value | reference/comment |
|---|---|---|
| $E_0(FA/FA^-)$ | -0.550 V | unchanged from [2, 19] |
| $E_0(Fd/Fd^-)$ | -0.430 V | unchanged from [2, 20] |
| $E_0(P700^+/P700)$ | 0.480 V | unchanged from [2, 21] |
| $E_0(NADP^+/NADPH)$ | -0.113 V | unchanged from [2, 22] |
